# Negative selection on complex traits limits genetic risk prediction accuracy between populations

**DOI:** 10.1101/721936

**Authors:** Arun Durvasula, Kirk E. Lohmueller

## Abstract

Accurate genetic risk prediction is a key goal for medical genetics and great progress has been made toward identifying individuals with extreme risk across several traits and diseases (Collins and Varmus, 2015). However, many of these studies are done in predominantly European populations (Bustamante et al., 2011; Popejoy and Fullerton, 2016). Although GWAS effect sizes correlate across ancestries (Wojcik et al., 2019), risk scores show substantial reductions in accuracy when applied to non-European populations (Kim et al., 2018; Martin et al., 2019; Scutari et al., 2016). We use simulations to show that human demographic history and negative selection on complex traits result in population specific genetic architectures. For traits under moderate negative selection, ~50% of the heritability can be accounted for by variants in Europe that are absent from Africa. We show that this directly leads to poor performance in risk prediction when using variants discovered in Europe to predict risk in African populations, especially in the tails of the risk distribution. To evaluate the impact of this effect in genomic data, we built a Bayesian model to stratify heritability between European-specific and shared variants and applied it to 43 traits and diseases in the UK Biobank. Across these phenotypes, we find ~50% of the heritability comes from European-specific variants, setting an upper bound on the accuracy of genetic risk prediction in non-European populations using effect sizes discovered in European populations. We conclude that genetic association studies need to include more diverse populations to enable to utility of genetic risk prediction in all populations.

---

The past decade of genome wide association studies (GWAS) has uncovered a plethora of trait associated loci scattered across the genome (The Wellcome Trust Case Control Consortium, 2007; The UK10K Consortium, 2015; Visscher et al., 2017; Bycroft et al., 2018). Geneticists have devoted many resources to turning these associations into risk prediction models that can be used to guide healthcare decisions for a variety of traits and diseases (Vilhjálmsson et al., 2015). Recent work has suggested these risk scores may be ready for clinical use (Khera et al., 2018, 2019). While individuals with high genetic risk for diseases have been found using these scores (for example atherosclerosis (Natarajan et al., 2017) and breast cancer (Maas et al., 2016)), challenges remain in applying these risk scores uniformly across populations. Recent analyses have suggested that because many of the largest studies are concentrated on European populations, risk scores may be biased and less informative in non-European populations (Scutari et al., 2016; Martin et al., 2017a; Kim et al., 2018; Martin et al., 2019; Mostafavi et al., 2019). There are several reasons why risk scores may not transfer well across populations. One possibility is that alleles have different effect sizes in different populations, owing to differences in interactions with the environment (Novembre and Barton, 2018). Another possibility is that differences in linkage disequilibrium (LD) between variants across populations means that causal variants may be tagged differently in non-European populations, leading to differences in effect sizes (Martin et al., 2017a; Wojcik et al., 2019). Finally, the original risk score performance in Europeans may be inflated due to stratification (Berg et al., 2019; Sohail et al., 2019).

Here we propose that an additional reason for the lack of transferability of risk scores is that each population has its own genetic architecture, owing to the evolutionary processes that give rise to traits. Under this reasoning, a population’s demographic history influences the number of causal variants and their frequencies, resulting in some phenotypic variance coming from causal variants that are population-specific. For example, work on the genetic architecture of skin color in African populations has uncovered distinct loci affecting the trait in each population, suggesting that populations with independent demographic histories can end up with different genetic architectures and causal variants for the same traits (Martin et al., 2017b). Indeed, modeling work suggests that genetic architecture is an outcome of the evolutionary process rather than a trait-specific property (Lohmueller, 2014).

Recent exponential growth in human populations has created an excess of new variants that tend to be low frequency and population specific (private variation; (Keinan and Clark, 2012; Tennessen et al., 2012)). Population genetic models of genetic architecture that include negative selection suggest that, in aggregate, low frequency variants could contribute substantially to traits (Eyre-Walker, 2010; Sanjak et al., 2017; Uricchio, 2019). Applications of these models to large-scale genetic datasets have discovered that many traits are under negative selection, ranging from anthropometric traits to molecular phenotypes (Hernandez et al., 2017; Gazal et al., 2017, 2018; Zeng et al., 2018; Schoech et al., 2019; Uricchio et al., 2019). Depending on the interplay between allele frequency and effect size, these variants could make up a large portion of the heritability for many traits, as demonstrated by a recent GWAS on height and BMI using whole genome sequencing data (Wainschtein et al., 2019). Because heritability is the proportion of variance explained by additive factors, it is directly related to the accuracy of phenotypic prediction as the variance explained by the genetic risk score (de los Campos et al., 2010). If these private variants do contribute to heritability, it follows that the variants will not be useful for risk prediction between populations because they are not present in other populations. The proportion of heritability that private variants explain places an upper bound on the accuracy of risk scores between populations.

In this study, we use simulations under demographic scenarios of recent population growth with varying amounts of negative selection as well as analyses of empirical data to test the role of private variants in complex traits.

We begin by conducting simulations under models of human evolution, where an ancestral population split into a group that underwent a genetic bottleneck out of Africa (representing a European population) and a group that stayed within Africa without a bottleneck (representing an African population; Fig 1a; (Gravel et al., 2011)), coupled with varying levels of negative selection on traits (including no negative selection). We include negative selection by modifying the relationship between a mutation’s effect on the trait and its effect on reproductive fitness using the model put forth by Eyre-Walker in 2010 (see Methods). This model includes a parameter, *τ*, which ties the selection coefficient of a mutation to its effect on a trait (see Methods; (Eyre-Walker, 2010)). Larger values of *τ* imply that more evolutionarily deleterious mutations have larger effects on the trait. We define private variants as those that occur in Europe and are absent from Africa and shared variants as those that occur in both populations. Note that by our definition, private variants may be shared between other Out-of-Africa populations due to shared recent history. Importantly, our model includes exponential growth in the out-of-Africa population, which creates an excess of private variants, as well as low levels of migration between the European and African populations, which can turn some private variants into shared variants. These simulations predict 11.98% of variants at a frequency greater than 5% in Europe will be private to Europe, suggesting that private variants need not have very low allele frequencies (Fig 1b). In the Exome Aggregation consortium (ExAC) dataset, we find that 11.69% of variants above 5% frequency in the non-Finnish European sample are private to Europe (see Supplementary note S3.3), suggesting the model predicts the amount of private variation accurately. In addition, the simulations predict there to be more causal variants ( |*s*| ≥ 1×10^−5^; Fig 1c; Methods) in Europe than in Africa, consistent with exponential growth resulting in many more variants in Europe.

**Fig. 1.**
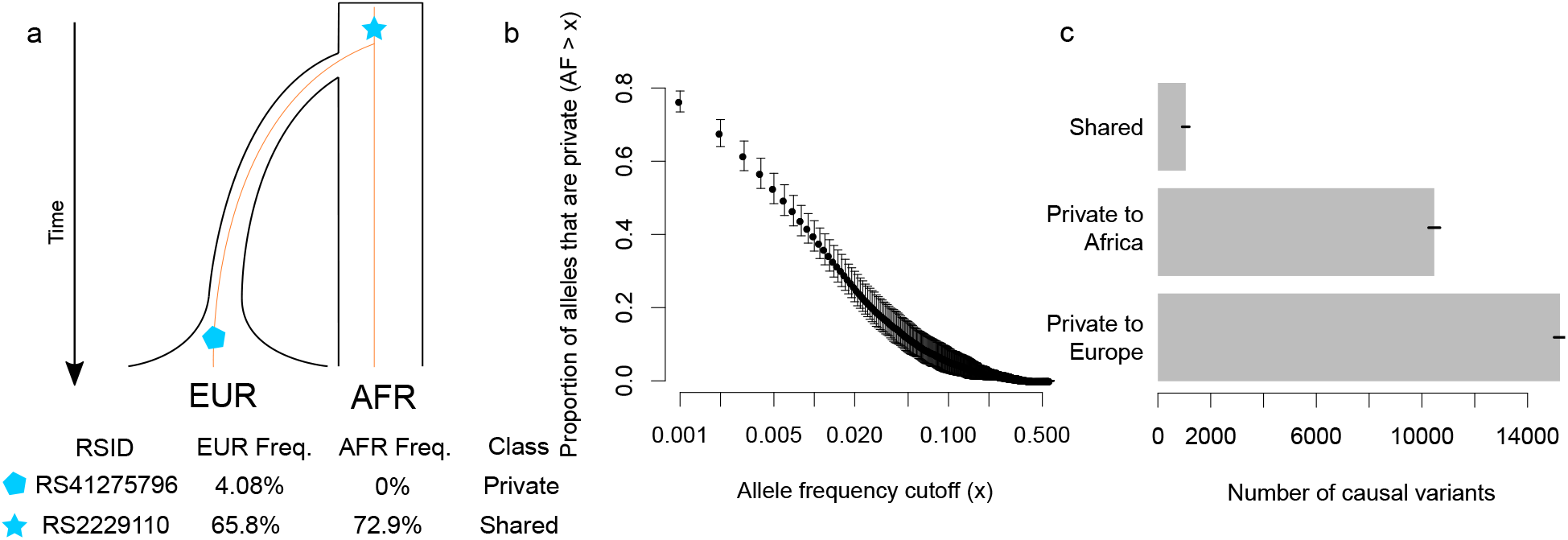
Human population history generates population-specific variants. a) model for variants that are shared (common to Europe [EUR] and Africa [AFR]) and private (occurring only in EUR and absent from AFR). Bottom: examples of private and shared variants from ExAC (Lek et al., 2016). b) the proportion of alleles at a frequency x or greater that are private to EUR (absent from AFR) in simulations from a realistic demographic model with negative selection. Note that the X axis is on a log scale. Error bars denote standard deviation over 10 replicates. c) the number of causal variants (defined as variants that affect fitness; |*s*| ≥ 1×10^−5^) for private and shared variants in simulated data under a realistic demographic history and negative selection. Lines indicate standard deviation over 10 simulation replicates.

We reasoned that since there are many private causal variants in our simulations, they may account for a substantial proportion of the heritability in aggregate. We examined the contribution of private variants to heritability and found that when traits are not tied to fitness (*τ* = 0), private variants account for ~30% of the heritability (Fig. 2a). However, when the coupling between trait effects and fitness effects is moderate (*τ* = 0.25) or strong (*τ* = 0.5), private variants account for over half of the heritability, with a maximum of ~79% under strong coupling (Fig. 2b,c). These results suggest that many causal variants, which jointly explain much of the heritability, tend to be population specific and this is a consequence of how the trait relates to fitness as well as the demographic history of the population.

**Fig. 2.**
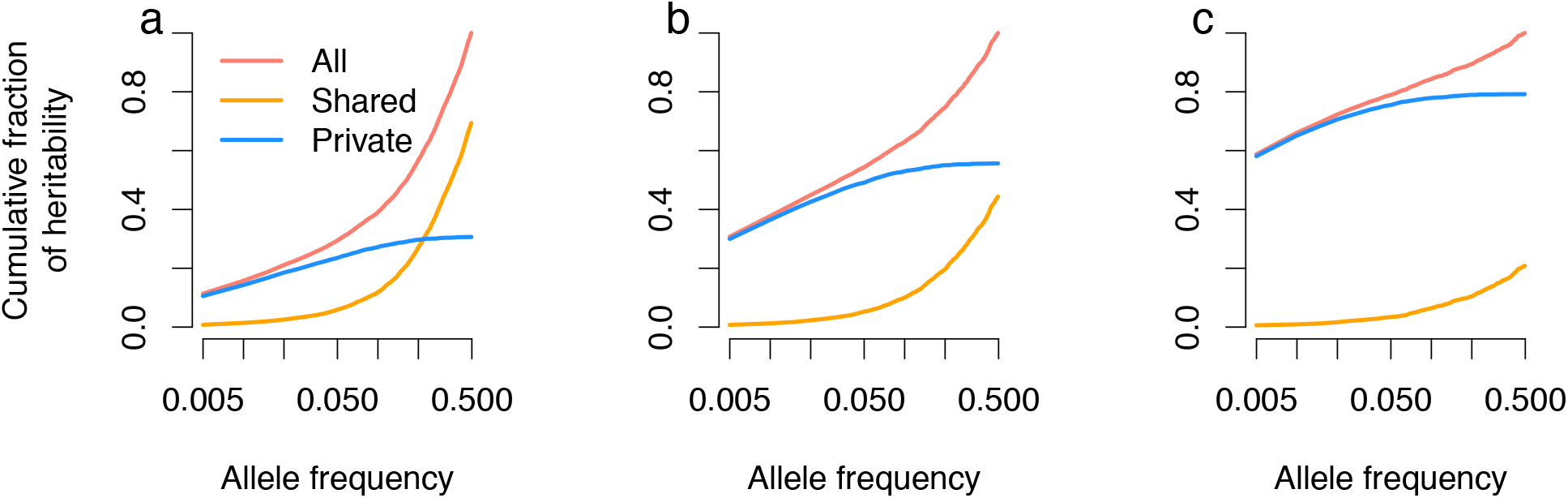
Cumulative fraction of heritability explained by private and shared variants under a) no relation between a mutation’s effect on fitness and the trait (*τ* = 0) b) moderate coupling between a mutation’s effect on fitness and the trait (*τ* = 0.25). c) strong coupling between a mutation’s effect on fitness and the trait (*τ* = 0.5). Note the X axis is on a log scale. As negative selection increases, a greater fraction of heritability comes from variation that is found only within Europe.

The fact that many of the variants that affect the trait are not shared across populations may limit the applicability of polygenic risk scores (PRS) derived from European populations to other populations. This effect would be distinct from imperfect tagging of causal variants due to differences in LD patterns between populations. To test for this effect in simulated data, we calculated true genetic risk scores for individuals in the simulated European and African populations and asked how well risk scores derived from only private variants and only shared variants correlated with the true risk scores. Risk scores derived from only shared variants represent the case where a polygenic risk score can be transferred from Europe to another population. If risk scores from shared variants correlate well with the true risk scores, genetic risk scores may still be accurate across populations. We note that these simulations include identification of the true causal SNPs and, as such, are much higher than PRS accuracies reported elsewhere (Martin et al., 2019). These simulations represent the best case scenario for risk scores. We found that when traits are not tied to fitness, the shared PRS has a 91% correlation in Europe and 96% correlation in Africa with the true PRS, suggesting that genetic risk scores can be applied between populations (Fig 3a). However, we found that when negative selection increases, the correlation between shared risk scores and the true genetic risk decreases (Fig 3bc) and the correlation between private risk scores and true genetic risk increases (Fig 3def). For traits with strong coupling between trait effects and fitness effects (*τ*=0.5), the correlation between the true risk scores and the risk scores derived from shared variants drops to 62% in Europe and 57% in Africa (Supplementary table S3). These findings suggest that genetic risk scores based solely on shared variants may be substantially less accurate than genetic risk scores using all variants and may not transfer between populations well when traits are under negative selection.

**Fig. 3.**
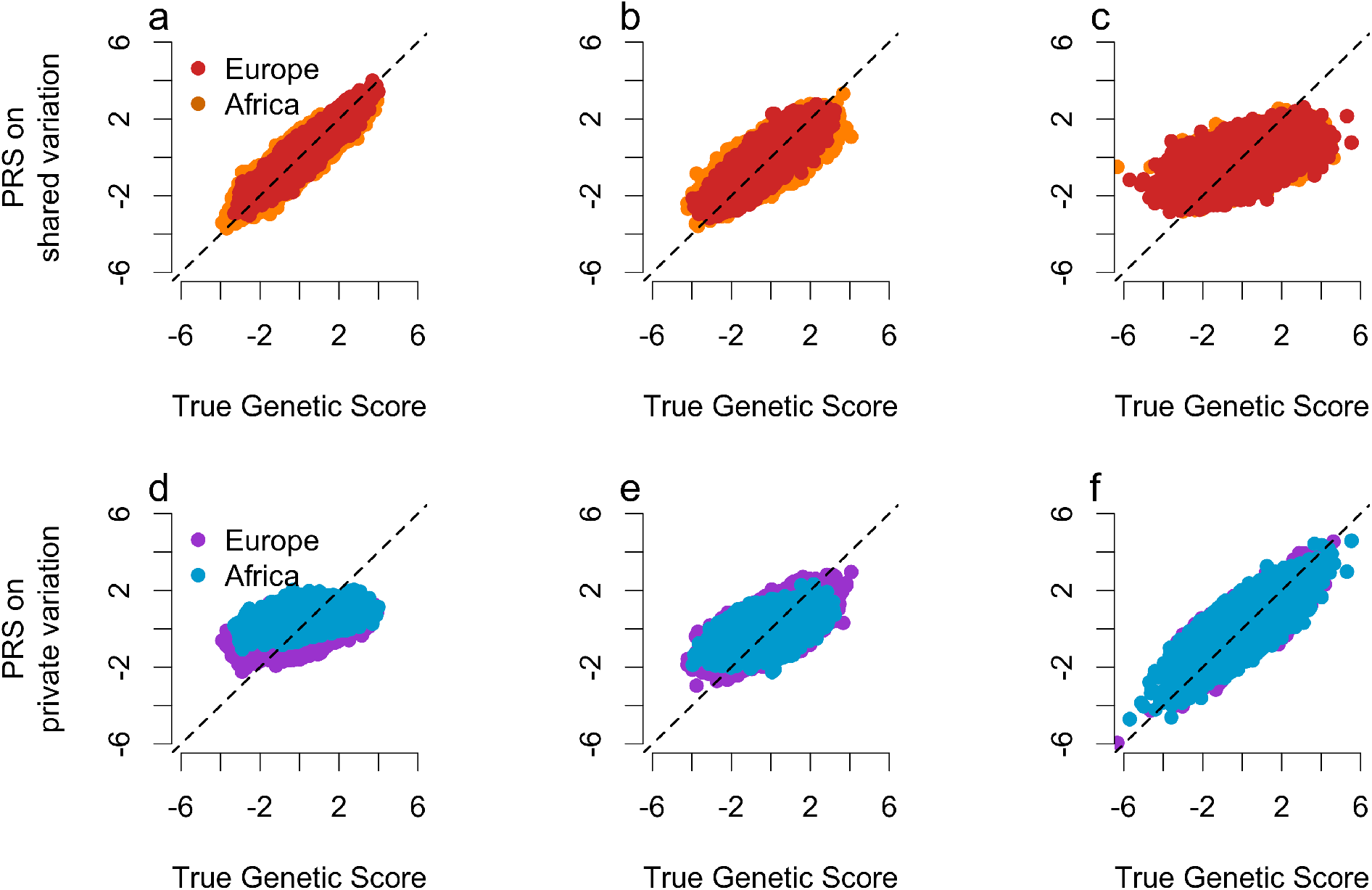
Genetic risk score accuracy for shared variants only (top row) and private variants only (bottom row) in Europe and Africa on simulated data with different degrees of purifying selection. The black line shows the 1:1 line. a,d) no relationship between a mutation’s effect on fitness and its effect on the trait (*τ* = 0) b,e) moderate coupling between fitness and trait effects (*τ* = 0.25) c, f) strong coupling (*τ* = 0.5). As the strength of coupling increases, risk scores computed from shared variation become less correlated with the true genetic risk score. However, at the same time, risk scores computed from private variation become more correlated with the true genetic risk scores.

While shared variants do not capture the full distribution of genetic risk for traits under negative selection, we asked whether individuals in the tail of the true risk distribution remained in the tail when examining shared variants only. When there is no coupling between fitness and trait effects (*τ*=0), we find that shared variants capture 35% of the tail correctly in Europe and 28% of the tail correctly in Africa (Table 1). However, when there is moderate coupling (*τ*=0.25), this number drops to 11% in Europe and 7% in Africa. When there is strong coupling, the risk score based on shared variants falls to 0% in both Europe and Africa, suggesting that shared variation cannot identify individuals at the highest risk for disease. In contrast, when considering only private variants for a trait under strong negative selection, the risk score correctly identifies 44-46% of individuals who are at the extremes of the risk distribution. These results suggest that when using scores derived from European populations, individuals who are truly in the tails of the risk distribution will not be identified using shared variants alone, corresponding to a high false negative error rate. In addition, the low recall for both of these risk scores suggests many individuals that are in the tails of the distribution will be missed.

**Table 1.** Percentage of individuals in the extreme 5% tail of the true genetic risk distribution that are recovered when using only private variants and shared variants in simulated European and African populations. Overall, the percentage of individuals correctly classified is low, suggesting that there will be many false negatives when using genetic risk scores to identify individuals in the tails of the risk distribution. Further, as the strength of selection increases, shared variants correctly classify fewer individuals, while private variants classify more individuals correctly.

| $\tau$ | Shared (Europe) | Private (Europe) | Shared (Africa) | Private (Africa) |
| --- | --- | --- | --- | --- |
| 0 | 35% | 22% | 28% | 11% |
| 0.25 | 11% | 20% | 7% | 18% |
| 0.5 | 0% | 46% | 0% | 44% |

While our simulations suggest private variants may be an important component of the heritability and may limit risk prediction across populations, their precise role depends on the extent of negative selection acting on traits, which remains an open question (Gazal et al., 2017, 2018; Hernandez et al., 2017; Simons et al., 2018; Schoech et al., 2019; Uricchio et al., 2019). Thus, we tested how much of the heritability private variants account for in real GWAS data in European populations, where GWAS data is abundant. We used summary statistics for 43 different traits and diseases from the UK Biobank relating to anthropometric and blood related traits as well as cancer-related and non-cancer related diseases (see URLs). We built a Bayesian model to classify variants as private or shared using the allele frequency conditional on a demographic model and distribution of fitness effects inferred for a European population (see Methods; Supplementary note S3). We evaluated the accuracy of our probabilistic framework by simulating data under the European demographic model and calculating the probability that each variant is private. We find the area under the receiver operator characteristic curve is 0.8 and at a false discovery rate (FDR) of ~5%, we have a recall of 98% (Supplementary fig. S1, Supplementary table S2). We evaluated the performance of our model on exome variants from ExAC (Lek et al., 2016) and found that 97% of the variants we predict to be private in the Non-Finnish European population (66,740 chromosomes sampled) were not observed in Africa (10,406 chromosomes from African and African American individuals) at a frequency greater than 5%. (Supplementary table 1). Together, this accuracy gives us confidence that our model is able to distinguish between private and shared variants based on allele frequency alone. Importantly, our determination of whether a variant is private or shared is expected to hold regardless of the sample size taken from either population (Supplementary note S3.2; Supplementary table 2).

Applying our model to real data, we find that the average SNP heritability explained by private variants is 50%, consistent with a moderate amount of selection on most traits (*τ* ≈ 0.2 − 0.3; Fig. 4). Examining categories of diseases, we find that cancer related diseases have 59% of the heritability in private variants, while non-cancer related diseases have 56% of the heritability in private variants. On the other hand, private variants account for ~40% of the heritability in blood related and anthropometric traits. These results are consistent with negative selection acting on a wide range of phenotypes and driving the heritability towards population specific variants. The fact that early onset and late onset diseases show similar patterns may be reflective of pleiotropic loci under similar selective pressures in both disease classes (Simons et al., 2018). To ensure that our results were not driven by shared SNPs that we mistakenly classified as private, we adjusted for both a simulation-based and empirically-based FDR. At the threshold used for classifying variants as being private, the simulations suggest the FDR is ~5% (Supplementary note S3.2) and empirical data suggest the FDR is ~14% (Supplementary note S3.3). We adjusted our estimates of the heritability attributable to private variants by removing 5% (and 14%) of the private SNPs that explain the most heritability (Figure 4). Despite the extremely conservative nature of this correction (because there is no a priori reason to suspect that the falsely inferred private SNPs should explain the most heritability), we find that a sizeable proportion of the heritability (about 20%) still comes from private variants. In addition, these estimates of the proportion of heritability explained by private variants likely underestimate the true proportion due to LD between tagging and causal variants (see Supplementary note section S2). Phenotypes with a majority of heritability explained by private variants are not likely to be predicted well in non-European populations, even if effect sizes are accurately inferred. Our analysis suggests that all traits examined here have greater than 30% of the heritability explained by private variants, indicating that cross population risk scores are limited in accuracy and many population specific causal variants remain to be discovered.

**Fig. 4.**
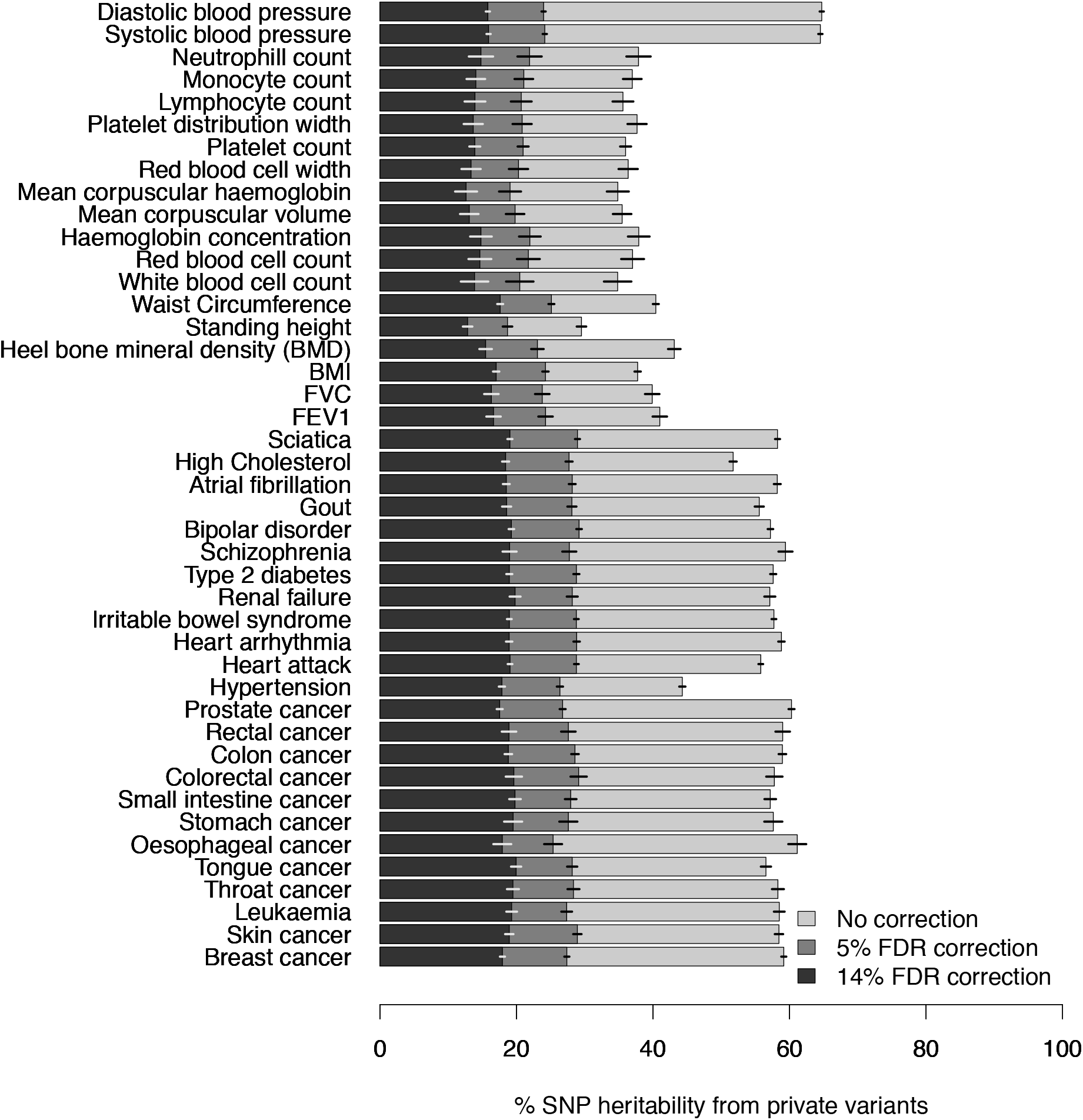
Percentage of heritability explained by private variants across 43 traits and diseases in the UK Biobank. Non-cancer and cancer diseases have the most variance explained by private variants, consistent with more negative selection on them versus blood and anthropometric traits. FDR corrected values come from removing 5% (darker grey) or 14% (black) of the private SNPs that explain the most of the variance, resulting in a conservative estimate of the heritability attributable to private SNPs. Lines indicate 95% confidence intervals obtained via 1 MB block jackknife.

In this work, we have shown that recent population growth and negative selection create population-specific genetic architectures for phenotypes, which has the direct effect of reducing the accuracy of genetic risk scores when applied between populations. The reduction in accuracy will depend on how differentiated populations are, with accuracy decreasing as populations before more differentiated. Another case to consider is admixed populations where some causal variants could be introduced and thus become shared variants. In these cases, we expect the utility of genetic risk scores to be higher, but this will depend on how recent the admixture was and how many causal variants are transferred between populations, which can vary between individuals.

We also highlight a crucial issue in identifying individuals in the tails of the risk distribution. If genetic risk scores are to be used more commonly, false negative rates must be more closely examined across populations and phenotypes. Our work suggests that many causal variants may not be shared between populations, indicating that variants ascertained in European populations may not be informative in other populations. This could occur because on average, more European specific variants have been either directly included in GWAS or imputed more often than variants specific to other non-European populations. To ensure equal predictive power of genetic risk scores across populations, whole genome sequencing based association studies must be undertaken in non-European populations. Such studies would allow for unbiased discovery of private variants accounting for much of the heritability, resulting in improved genetic risk prediction in non-European populations. Finally, large imputation panels from the relevant population of interest are necessary to include variation not present in Europe.

## Methods

### Population genetic modelling and simulations

We performed forward simulations using SLiM v3 (Haller and Messer, 2017). We simulated a demographic history for a European and an African population according to the demographic model fit by Gravel et al (including migration). We simulated a mutational target size of 5 megabases with a mutation rate of 1.2×10^−8^ and a recombination rate of 1×10^−8^. For new mutations, we draw selection coefficients from a gamma distribution with parameters fit by Kim et al (mean −0.01026, *α*=0.186). We sample 10,000 haploid genomes from each population. To simulate a quantitative trait, we follow the model described by Eyre-Walker and the framework set by Lohmueller (2014) where a SNP’s effect on a trait, *β*, is given by

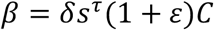

where *δ* ∈ {−1, 1} with equal probability, *ε*~*N*(0, 0.5), and *s* is the selection coefficient of a variant segregating in the population at the end of the simulation. *C* is a scaling factor for effect sizes and controls the heritability for a given mutational target size. In these simulations, *C* was set to obtain a heritability of ~0.4 (see Supplemental table S4). Finally, *τ* reflects the relationship between a SNP’s effects on fitness and the trait. *τ* = 0 indicates no relationship between fitness and the trait, while *τ* > 0 suggests that negative selection acts on the trait, with larger values indicating stronger negative selection. We call variants private and shared based on their allele frequency in a sample of 10,000 chromosomes from both populations (see below).

### Risk Score Calculation

We compute 3 sets of risk scores: 1) using all variants 2) using variants private to the simulated European population 3) using variants shared between the simulated European and African populations. For each haploid genome, we sum the effect sizes *β* for each class of variants, resulting in 3 scores for each genome. We standardize the scores by subtracting the mean of the true genetic risk (class 1) and dividing by the standard deviation of the true genetic risk (class 1). We compute the Pearson correlation between classes 1 and 2 as well as 1 and 3.

### Model to identify private variants

When analyzing the empirical UK Biobank data, it is challenging to assess whether a particular variant is private or shared. If a variant is seen only in one population, it is possible that it is truly private to that population, or instead, it is shared, but at too low a frequency to have been discovered with the number of individuals samples from that other population. To address this issue, we built a probabilistic model to evaluate the probability that a variant is private to a population given the number of copies of the allele in that population (that is, the allele frequency). Given a collection of variants, we wish to compute this probability for each variant *j*:

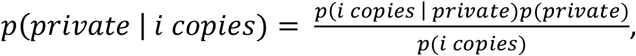

where *p*(*private*) is 1 if the variant is private to a population and 0 if it is shared. The quantity *p*(*i copies*) is the probability of an allele having *i* copies in the population and can be computed from the site frequency spectrum. The quantity *p*(*i copies* | *private*) represents the probability of a variant having *i* copies in the population, given that it is private and can be computed from the site frequency spectrum of private variants rather than the spectrum of all variants. While these two quantities can be computed analytically (see Wakeley and Hey, 1997; Wakeley, 2008), here we are concerned with non-equilibrium demographies and a non-neutral distribution of fitness effects. Thus, we instead obtain these quantities from forward simulations. Using the simulation framework described in Supplementary note section S3.1, we compute the SFS for private variants (giving us *p*(*i copies* | *private*)), the SFS for all variants (giving us *p*(*i copies*)), and the overall probability that a variant is private (giving us *p*(*private*)). Importantly, these simulations do not make any assumptions about the relationship between a mutation’s effect on the trait and its effect on fitness (i.e. *τ*). Rather, we only use the simulations to quantify how the demography and purifying selection (irrespective of the trait) will affect the SFS.

### Model evaluation

We evaluated the ability of our model to distinguish between private and shared variants by simulating new data and performing binary classification, calling a variant private if the *p*(*private* | *i copies*) exceeded some threshold *t*. We varied this threshold and computed the number of true positive (private variants that are truly private), false positives (private variants that are truly shared), false negatives (shared variants that are truly private), and true negatives (shared variants that are truly shared). We summarized this using receiver operator characteristic and precision recall curves (Supplementary note S3.2; Supplementary figure S1; Supplementary tables S1, S2).

We also validated our model using data from the Exome Aggregation Consortium (Lek et al., 2016). For each variant in ExAC, we used our model to compute the probability that the variant is private to the Non-Finnish European population based on the allele frequency in that population. Then, we checked whether variants were observed in a sample of 10,406 African and African-American samples (Supplementary note section S3.3).

### Partitioning *V*_*A*_

We applied our model to GWAS summary statistics from 43 traits in the UK Biobank released by the Neale lab (see URLs). We used our model to compute the probability that each variant is private. We computed the additive genetic variance for variants with a high posterior probability of being private to the British cohort and divided that by the total amount of additive genetic variance explained by SNPs (Supplementary note section S4). We also performed the inference using a randomized algorithm to correct for the effects of linkage disequilibrium and misestimated effect sizes as well as population stratification (Supplementary note section S2, S4; Supplementary figs S2, S3, S4, S5, S6, S7, S8). Finally, we also independently replicated the results on BMI using data from the GIANT consortium (Supplementary note section S4).

### Correcting for falsely inferred private variants

The proportion of heritability from private variants depends on the threshold used to classify variants as private or not. In Figure 4, we use a threshold that corresponds to an estimated false discovery rate (FDR) of ~5% from simulations and ~14% from empirical data (Supplementary note S3). We also conservatively estimate the proportion of the heritability attributable to private variants accounting for these falsely inferred private SNPs (Figure 4). To do this, we sorted the private SNPs in decreasing order of the amount of variance that the explained and removed the top 5% of these SNPs (i.e. we removed the 5% of the private SNPs that explained the most heritability). We also repeated this procedure removing 14% of the SNPs. This filter corresponds to a worst-case scenario where the falsely inferred private SNPs contribute the most to heritability.

## Acknowledgements

We thank Sriram Sankararaman, James Boocock, Alec Chiu, and Ruth Johnson for helpful discussions and Bogdan Pasaniuc, Nelsen Freimer, and members of the Lohmueller lab for helpful comments on a draft of this manuscript. AD is funded by NSF Graduate Research Fellowship DGE-1650604 and KEL is funded by NIH Grant R35GM119856.

## URLs

GWAS summary statistics http://www.nealelab.is/uk-biobank

