## Supplementary Information for "Negative selection on complex traits limits genetic risk prediction accuracy between populations"

### Supplementary material for 'Negative selection on complex traits limits genetic risk prediction accuracy between populations'

Arun Durvasula and Kirk E. Lohmueller

**S4 Application to GWAS summary statistic data** 9

#### S1 Defining the proportion of heritability from private variants

Here we define how we can conceptualize the portion of the heritability that comes from variants that only occur in one population (i.e. private variants). We begin by describing a model in which an individual $i$ in a population $\phi$ has a phenotype $y_i$ that is a linear combination of genotypes ($\mathbf{x_i}, x_{ij} \in \{x_{i1}, ..., x_{iM}\}$), effect sizes ($\mathbf{b}, b_{ij} \in \{b_{i1}, ..., b_{iM}\}$), and a normally distributed term describing the effect of the environment $e_i \sim N(0, V_e)$:

$$y_i = \mathbf{x_i^T b} + e_i$$

The narrow sense heritability, $h^2$, of the phenotype $\mathbf{y}$ in the population is given by

$$h^2 = \frac{V_A}{Var(\mathbf{y})}$$

where the variance of the phenotype can be decomposed into additive, dominance, interacting, and environmental terms: $Var(\mathbf{y}) = V_A + V_D + V_I + V_E$. The additive genetic variance is $V_A = 2\sum_{j=1}^{M} p_j(1-p_j)b_j^2$ when there are $M$ variants and where $p_j$ is the allele frequency for variant $j$ and $b_j$ is the effect size of variant $j$.

We wish to examine the proportion of heritability that comes a particular class of variants. Consider a sister population $\psi$ that diverged from the population described above ($\phi$). Variants in population $\phi$ can be partitioned into those that appear only in $\phi$ (private variants) or those that appear in both populations (shared variants). The total number of variants is the sum of the number of shared and number of private variants, $M = M_p + M_s$. We wish to partition the heritability into these two classes, $h_p^2$ and $h_s^2$, which make up the total heritability: $h^2 = h_p^2 + h_s^2$. Define $\rho$ to be the the proportion of the heritability accounted for by the private variants.

The quantity of interest, then, is

$$\rho = \frac{h_p^2}{h^2} = \frac{V_{A,p}}{V_A}$$

The additive genetic variance from private variants is $V_{A,p} = 2\sum_{j=1}^{M} p_j(1-p_j)b_j^2 z_j$, where $z_j$ is an indicator function that is 1 when the variant $j$ is private to the population and 0 otherwise. We wish to obtain an estimator of $\rho$, $\hat{\rho}$.

### S2 The effect of linkage disequilibrium on our inferences

Our inferences of the heritability due to private variants made the assumption that the estimated effect sizes for the GWAS SNPs were the true effect sizes of the causal variants. Further, we assumed that the variants were all independent of each other. In truth, these assumptions are violated for a variety of reasons. First, due to linkage disequilibrium (LD), SNPs may be correlated with one another. Second, some of the non-zero effect sizes of GWAS SNPs may be due to the fact that the GWAS SNP is tagging (in LD with) an untyped causal variant and is itself not causal. Third, even if the GWAS variants analyzed in our study are the true causal variants, their effect sizes may be mis-estimated by the effects at nearby SNPs in LD with them.

Thus, given these challenges, we carefully considered the effect that unmodeled LD may have on our inferences.

In principle, recent methods such as stratified LD score regression (S-LDSC; [3]), SumHer [17], or HESS [14] could provide us with the ability to examine the proportion of heritability that comes from private variants. These methods attempt to deconvolve the observed marginal association statistics, $\hat{b}_j$, by modeling linkage disequilibrium (LD) between tagging variants and causal variants. However, these methods rely on out of sample estimates of the amount of LD between variants (*i.e.* from a reference population). While estimates of LD (measured by $\hat{r}^2$) between common variants can be obtained with minimal variance in some cases, this depends greatly on allele frequency, as outlined below.

Consider two loci, $A$ and $B$, with allele frequencies $p_A$ and $p_B$ of the minor allele. The correlation coefficient, $r^2$, is defined as:

$$r^2 = \frac{(p_{AB} - p_A p_B)^2}{p_A(1-p_A)p_B(1-p_B)}$$

An estimator of $r^2$ is obtained by using estimates of $\hat{p}_{AB}$, $\hat{p}_A$, and $\hat{p}_B$ [15, 8]. If the allele frequencies are assumed to be binomially distributed and if the sample size used to obtain estimates of allele frequencies is large enough, we can model these frequencies as approximately normally distributed. Then, the variance of the estimator, $Var(\hat{r}^2) \approx (1-r^2)^2$ [4]. This relationship suggests that when the correlation is lowest, the variance will be the largest. Analysis of the mathematical properties of $r^2$ as a measure of LD suggests that the maximum value of $r^2$ is lowest when $p_A$ is much different from $p_B$ [19]. Taken together, this suggests that there will be much uncertainty when using LD panels to estimate LD between common and rare variants.

Here, we are interested in LD patterns between private variants, which tend to be rare, and shared variants, which tend to be common. Therefore, in lieu of using previous methods that rely on out of sample LD estimates, we consider several approaches that do not rely on knowing the patterns of LD between variants: 1) We repeated our inference only considering a sub-set of SNPs that would have a greater chance of being independent of each other, 2) We used coalescent simulations to investigate the effect of unaccounted for LD on the inference of the heritability contained in private variants, and 3) We tested whether the effect sizes of private variants or shared variants were more likely to be over-estimated due to LD. All of these analyses suggest that while unmodeled LD between SNPs may bias some of our inferences, they are biased in the direction of underestimating the heritability due to private variants. In other words, our estimates are likely to be conservative and our overall conclusion that a substantial amount of the heritability for anthropometnric and disease traits is due to private variants is robust to LD. Below we detail these results of these three analyses.

#### S2.1 Inference of $\rho$ by pruning for LD

Here, we describe an estimator for proportion of heritability that comes from private variants when using a sub-set of SNPs within a genomic window. By thinning the SNPs, we should remove some of the effects of LD between the GWAS SNPs. Our estimator has the form:

$$\hat{\rho} = \frac{\hat{V}_{A,p}}{\hat{V}_A}.$$

First, we describe a procedure for estimating $\hat{V}_{A,p}$. Consider a window $w$ with $K$ SNPs. We randomly select 1 SNP, $k$, and compute the additive genetic variance for that SNP:

$$\hat{V}_{A,p,w,k} = 2\hat{p_k}(1-\hat{p}_k)\hat{b}_k^2 z_k$$

where $z_k$ is an indicator function that equals 1 when $p(\omega|\hat{p}_k) \geq t$ (see Section S3 for a description of this model) and 0 otherwise. We repeat the procedure $T$ times and take the sum:

$$\hat{V}_{A,p,w} = \sum_k^T \hat{V}_{A,p,w,k}$$

Then, to estimate $\hat{V}_{A,p}$, we sum over the windows:

$$\hat{V}_{A,p} = \sum_w \hat{V}_{A,p,w}$$

If $T$ is large and $w$ contains a single causal variant, then $E[\hat{V}_{A,p,w}] \approx 2p_c(1-p_c)(b_c\bar{r}_c^2)^2$, where $\bar{r}_c^2$ is the average LD of variants in the window and the causal variant[1].

A similar estimator for $\hat{V}_A$ can be obtained:

$$\hat{V}_{A,w} = 2\sum_{k}^{T} \hat{p_k}(1-\hat{p}_k)\hat{b}_k^2$$

$$\hat{V}_A = \sum_{w}^{W} \hat{V}_{A,w}$$

We obtain standard errors by using a jackknife over the windows [12]. We denote the estimate of $\rho$ obtained using the full data set (*i.e.* all windows) as $\hat{\rho}$. Further, let $\hat{\rho}_{(-w)}$ be the estimate of $\rho$ obtained by deleting window $w$ and let $\bar{\rho} = \frac{1}{W}\sum_{w=1}^{W}\hat{\rho}_{(-w)}$. We compute the standard error ($SE_\rho$) and bias ($B_\rho$) as

$$SE_\rho = \sqrt{\frac{w-1}{w}\sum_{w=1}^{W}(\hat{\rho}_w - \bar{\rho})^2}$$

$$B_\rho = (w-1)(\bar{\rho} - \hat{\rho})$$

We obtain 95% confidence intervals by first bias correcting the estimate ($\hat{\rho}_{corr} = \hat{\rho} - B_\rho$). Then, the confidence interval is $\hat{\rho}_{corr} \pm z(\alpha)SE_\rho$, where $\alpha$ is the confidence level, $z(\alpha) = \sqrt{2}erf^{-1}(\alpha)$ and $erf^{-1}(x)$ is the inverse error function. In all plots using this estimator, we plot the bias corrected estimate rather than the full estimate.

This approach to confidence intervals ignores correlations between windows, which could lead to double counting variants if causal variants are clustered together. We found that when we varied the window size between 500 SNPs ($\approx$ 100KB), 5,000 SNPs ($\approx$ 1MB), and 50,000 SNPs ($\approx$ 10MB), the results were consistent, suggesting that this was not a practical issue (Section S4).

From the expectation above, it is clear that the estimator we propose is biased, $E[\hat{\rho}] \leq \rho$. Next, we show that this bias conservatively estimates the proportion of heritability from private variants.

When $\bar{r}_c \approx 1$, the average LD in the window between the tagging variants and causal variants is close to 1, resulting in $E[\hat{\rho}] \approx \rho$. However, such tightly linked regions will not be common in the human genome. Several papers have noted that the range of $r^2$ depends on the allele frequencies of the loci in question (VanLiere and Rosenberg 2008). Specifically, as the two allele frequencies diverge, the maximum $r^2$ value decreases. In our case, shared variants tend to have larger allele frequencies than private variants, suggesting that the maximum allele frequency difference will occur when the tagging variant is private and the causal variant is shared or when the tagging variant is shared and the causal variant is private.

In the case where there is a mismatch between the categories of the tagging and causal variants, it is possible to end up with variation contributing to the wrong additive genetic variance bin. For example, if the tagging variant is shared and the causal variant is private,

[1] We do not know the true allele frequency of the causal variant ($p_c$), so the expectation of the estimator is not exactly equal to the quantity on the right hand side. However, if $r^2$ is large, then $p_c \approx p_t$ where $p_t$ is the allele frequency of the tagging variant and the expectation will hold.

$V_A$ that should be in the private bin would be apportioned into the shared bin, leading to downward bias in the estimate of $\rho$. On the other hand, if the tagging variant is private and the causal variant is shared, $V_A$ that should be in the shared bin will be in the private bin, leading to overestimates of $\rho$. However, in both of these cases, because of the dependency of $r^2$ on allele frequency, we do not expect these cases to contribute much to $V_A$ because $r^2$ will be low between the tagging and causal variants. Further, we expect the second case (where causal variants are shared and the tagging variants are private) to be uncommon when a trait is under negative selection because negative selection will push large effect alleles to lower allele frequencies, increasing the probability that these alleles are private. Thus, the scenario under which our estimate of $\rho$ would be upwardly biased are likely to be unusual.

On the other hand, when both the tagging and causal variant are in the same class (*i.e.* both private or both shared), the frequencies will be more similar, leading to estimates of $V_A$ that are reduced by a factor related to the amount of LD between the tag and the causal variant.

#### S2.2 Coalescent simulations to assess the performance of the estimator of $\rho$

We tested our intuition in Section S2.1 by performing coalescent simulations. We simulated genetic variation data using msprime [10] under the demography from Gutenkunst et al [6]. We simulated a single window and randomly assigned a causal variant within that window with an effect size $b \sim N(0,1)$. Then, for each of the non-causal variants in the window, we simulated a GWAS by assigning marginal association statistics as $\hat{b} = r^2 b$, where $r^2$ is the LD between a given variant and the causal variant.

Then, to simulate our estimation procedure (Section S2.1), we randomly pick a SNP from the window, and compute the additive genetic variance using the 'marginal' association statistic ($\hat{V}_A = 2p(1-p)\hat{b}^2$). We estimate $\hat{\rho}$ using the allele frequency to infer if an allele is private (see Section S3). We obtain the true additive genetic variance using the true effect size ($b$) and determine if an allele is private based on whether it is observed in the simulated African population. We repeat this procedure 10,000 times for each window and simulate 10,000 windows.

Then, to ensure our simulation is close to our real inference procedure, we compute $E[\hat{\rho}]$ by inferring whether alleles are private or shared (rather than using the true status from the simulation). This is achieved using the framework described in Section S3. In Figure S2b, we plot the estimated additive genetic variance from each of the windows versus the true additive genetic variance from each of the windows for private variants. As expected, we see that there is a downward bias, consistent with the effects of LD on $\hat{b}$. This means that LD causes us to underestimate the heritability accounted for by private variants. Finally, we compare $E[\hat{\rho}]$ to $\rho$ (Figure S2a) and see that the estimator is downwardly biased as well, consistent with a greater effect of bias on private variants compared to shared variants. Therefore, we conclude that the estimator described in Section S2.1 is conservative with respect to the amount of heritability explained by private variants.

#### S2.3 The relationship between marginal effect size estimates and LD

We wondered whether our estimates of $\rho$ could be biased by differences in tagging causal variation between private and shared variation. Because shared variants tend to be more common, they will tend to be in LD with more (and therefore tag more) variants. If this is the case, then shared variants will have inflated marginal effect sizes compared to private variants. This would lead to overestimating the additive genetic variance from shared variants compared to private variants, again making our inferences conservative.

To test this for this effect, we examined the summary statistics from a GWAS on BMI in the UK Biobank and correlated the marginal effect sizes with recombination rates estimated from patterns of LD in Europeans [9] for both classes of variants. We found that variants we predict to be private have a lower correlation (Spearman's $\rho = -0.04$, $P < 2 \times 10^{-16}$) between marginal effect sizes and recombination rates compared with variants we predict to be shared (Spearman's $\rho = -0.08$, $P < 2 \times 10^{-16}$). Said another way, the relationship between effect size and LD (which is one of the main ideas behind LD score regression; [2]), is stronger for shared variants than for private variants. This finding suggests that estimates of additive genetic variance from private variants are downwardly biased compared to estimates from shared variants, consistent with the intuition from Section S2.1 and results from our simulations (Section S2.2).

### S3 Detecting private variation

All of our inferences require accurately determining which variants are private to a population and which are shared. In principle, this is an empirical question. If enough genomes are sampled from two populations, it is possible to determine exactly which variants do not appear the other population. However, in practice, genomes are not evenly sampled from populations and differences in coverage, read mapping, and genotype quality can lead to biased empirical estimates of private alleles. Further, sample sizes may not be adequate to confidently assess that a particular variant it not polymorphic in a given population. Instead, it could be at low-frequency in that population and not detected in the sample. Thus, we develop a probabilistic approach to infer the probability that a variant is private, given its allele frequency. We then validate our approach using both simulated and empirical data.

#### S3.1 Probabilistic model to infer whether a variant is private

Here, we describe a probabilistic model to determine whether an allele is private or shared. We begin with the intuition that rare alleles tend to private and common alleles tend to be shared between populations, even in the presence of migration (migration can be thought of as sampling alleles from one population and placing them in the other population. Under this model, rare alleles will tend to stay within a population and not transfer between populations). This suggests that allele frequency is informative in determining whether an allele is private or not.

In Wakeley and Hey [20], the authors use coalescent theory to determine the frequency spectrum of private variants. We build on that theory to calculate the following probability:

$$P(\omega|i) = \frac{P(i|\omega)P(\omega)}{P(i)}$$

where $i \in \{1, ..., n\}$ is the number of copies of the allele in the sample ($n$) and $\omega \in \{0, 1\}$ is 1 if the allele is private and 0 if not. $P(i|\omega)$ is given by the site frequency spectrum of private variants, and $P(i)$ is given by the full site frequency spectrum. For example, in a constant sized equilibrium population, $P(i) = (\theta/i)/(\sum_i \theta/i)$. $P(\omega)$ is the prior probability of a variant being private to a population.

Wakeley and Hey [20] provide expressions to obtain these quantities in a constant sized equilibrium population without natural selection. However, here we are concerned with populations that are not in equilibrium and with variants under negative selection, so we obtain these probabilities via simulation under a particular demographic model $\mathcal{M}$ and distribution of fitness effects $\Gamma$.

In the results presented here, we use the demographic model from Gravel et al [5] that relates European and African populations. We use a distribution of fitness effects from Kim et al [11], assuming that mutations are additive (that is, $h = 0.5$) and that selection coefficients, $s$, are drawn from a gamma distribution with $\mu = -0.01026$ and $\alpha = 0.186$. Using these parameters, we simulate data for 10,000 European chromosomes using SLiM [7] and compute 1) the proportional site frequency spectrum for private variants ($P(i|\omega)$), 2) the proportional site frequency spectrum for all variants ($P(i)$), and 3) the proportion of private variants ($P(\omega)$). We defined private variants in the simulation as those that appear in the simulated European population but not the simulated African population.

Next, we store these quantities in a lookup table and use them to compute the probability that a variant is private given the number of copies of the allele in the empirical data. In the UK Biobank dataset, alleles are present at frequency $1 \times 10^{-6}$ and higher. However, in simulations, the lowest allele frequency is $1 \times 10^{-4}$. For alleles below this frequency, we set the probability equal to the probability for alleles at 1 in 10,000 (*i.e.* $P(\omega|i \leq 1/1 \times 10^{-4}) = P(\omega|1/1 \times 10^{-4})$).

#### S3.2 Simulation-based evaluation of the performance of our probabilistic model

We evaluated the ability of our model to distinguish private from shared variants using a combination of simulations and empirical data. First, we simulated a new dataset under the same demographic model and recorded whether each allele was observed in both populations. Then, we calculated the probability of each allele being private to the European population. We classified variants as private if the probability, $P(\omega|i) \geq t$, where $t$ is some probability cutoff. For each cutoff, we calculated 1) the number of variants that we predict are private and are truly private (true positives) 2) the number of variants that we predict are private and are truly not private (false positives) 3) the number of variants that we predict are not private and are truly private (false negatives) 4) the number of variants that we predict are not private and are truly not private (true negatives).

We summarize these numbers using two curves, a precision-recall curve (Figure S1a) and a receiver operator characteristic curve (ROC) (Figure S1b). We find that at a precision of 94%, we have a recall of 99% and that the area under the ROC curve is 0.80, suggesting that our model is able to distinguish between private and shared variants based on allele frequency alone (Table S2). We also tested the model on a simulated dataset including five times more individuals than the 10,000 individuals used in the initial simulation. Importantly, for this comparison, we used the same lookup table, based on 10,000 individuals, as before. This allows us to test how sample size affects our inferneces. We find that the precision-recall curve is largely the same, but there is a decrease in the ROC curve (AUROC= 0.70).

In addition, we plot $P(\omega|i)$ versus the allele frequency in the simulated independent data set (Figure S1c). We find that alleles higher than $\sim 10\%$ frequency have a negligible probability of being private, consistent with the intuition that common alleles are unlikely to be private.

We also examined several posterior probability thresholds in detail (0.1, 0.23, and 0.4; Table S2). Across these thresholds, we find that the FDR from simulations is $\approx 5\%$, suggesting that the model is relatively robust to the threshold used.

#### S3.3 Empirical evaluation of the performance of our probabilistic model

Next, we empirically validated the performance of our model to infer whether variants are private. Using data from the Exome Aggregation Consortium (ExAC; [13]), we use our framework described above to calculate the probability that each variant is private using the allele frequency in the Non-Finnish Europeans (NFE). In Figure S1d, we plot this probability for a random subset of 10,000 variants. We see that variants above 10% frequency have a very low probability of being private and that variants below that frequency increase in probability as the frequency decreases.

In addition, we classified variants in ExAC as private to EUR using a false discovery rate of 5% ($P(\omega|i) \geq 0.23$) and checked whether those variants were present in the 'AFR' subset of samples (Table S1). We see that 83% of the variants we call private are not observed in 'AFR' in a sample of 10,406 chromosomes. However, the 'AFR' sample in ExAC is a mixture of African American and African samples. Importantly, African-American samples are admixed between European and African populations [13]. This has the effect of introducing European variants into the 'AFR' samples, making variants we expect to be private to EUR appear shared.

To mitigate this problem, we introduce an allele frequency filter in the 'AFR' since we would expect these introduced variants to be at a low frequency in this mixture of African-American and African samples. We used filters of 0.001, 0.005, 0.01, and 0.05. We see that as the filters move higher in frequency, the fraction of alleles that are predicted private and not observed in 'AFR' increases, reaching 0.97 with a filter of 0.05.

An alternate approach to control for admixture is to use the observed allele frequency in the African American sample ($f_{AA}$), the expected admixture fraction ($\alpha$), and the observed allele frequency in the European sample ($f_{EUR}$) to estimate the true African allele frequency ($f_{AFR}$).

In particular, if the admixture fraction from Europeans intro Africans is approximately 20% ($\alpha \approx 20\%$), then the frequency of the European allele in the African American sample will be $f_{AA} = (1-\alpha)f_{AFR} + \alpha f_{EUR}$. An estimate of $f_{AFR}$ can be obtained by rearranging and plugging in the maximum likelihood estimates of the allele frequency:

$$\hat{f}_{AFR} = \frac{\hat{f}_{AA} - \alpha \hat{f}_{EUR}}{(1-\alpha)}$$

We use this to estimate the frequency of each allele in the AFR population. Next, for each allele, we simulate drawing a sample of 50,000 chromosomes from the AFR population by sampling from a binomial distribution:

$$E[X_{AFR}] \sim Binom(50000, \hat{f_{AFR}})$$

where $X_{AFR}$ is the count of the allele in the AFR population. Then, for each variant, we call the true status of the allele private if the expected count is 0 ($X_{AFR} = 0$).

We compute the overall empirical FDR by counting how many alleles are predicted private by the model and are truly private (true positives) and by counting the number of alleles that are predicted private by the model and are not private (false positives). We summarize the results across 3 thresholds in Table S2. We find that the empirical FDR is $\approx 14\%$ across these thresholds.

Taken together, these simulations and empirical evaluations suggest that our model is able to distinguish between private and shared variants. Additionally, out of an abundance of caution, we will utilize the simulation-based FDR of 5% and the empirical-based FDR of 14% in downstream inferences (Section S4).

### S4 Application to GWAS summary statistic data

We applied our partitioning method to summary statistic GWAS data computed on individuals from the UK Biobank. We used summary statistics for 43 traits and diseases released by the Neale lab at `http://www.nealelab.is/uk-biobank/`. Details of the procedure for GWAS, including quality control can be found at the website, but we briefly summarize it here.

For sample inclusion, the authors used unrelated samples and subset to British individuals using PCA and self-reported ancestry ('white-British'/'Irish'/'White'). This resulted in 361,194 samples. SNPs were imputed using the Haplotype Reference Consortium, the UK10K project, and the 1000 Genomes data. Variants were retained if they had an INFO score $> 0.8$, minor allele frequency $> 0.0001$ (MAF $> 1 \times 10^{-6}$ if annotated as missense or protein truncating by VEP), and a Hardy-Weinberg equilibrium $p$-value $> 1 \times 10^{-10}$. This filter resulted in 13.7 million SNPs. The association testing was done using the first 20 principle components, sex, age, as well as interaction and non-linear terms for sex and age. We used summary statistics for GWAS performed including both sexes and for quantitative phenotypes, we used inverse rank normalized phenotypes (as opposed to the raw phenotypes).

From the summary statistic data, we use the minor allele frequency in the British cohort and the effect size to compute the additive genetic variance (Section S2.1) and the probability that the variant is private to Europe (Section S3). In the main text, we report results using all SNPs reported in the summary statistics. These results use a probability threshold of $P(\omega|i) = 0.23$, corresponding to a simulated FDR of $\approx 5\%$ and an empirical FDR of $\approx 14\%$. In the main text, we also present results using the FDRs to correct for the fact that some of the SNPs we infer to be private are likely false (i.e. they are truly shared). Specifically, the FDRs in Table S2 suggest that $\approx$5-14% of the SNPs we call private are actually shared. We performed a conservative correction by removing the top 5% and 14% of SNPs based on the amount of heritability they explain and plot the results from the remaining SNPs. This is a conservative correction (i.e. the final results end up under-estimating the amount of the heritability attributable to private variants) because this correction assumes all the SNPs we misclassify explain the top 5% and 14% of genetic variance.

We also show the results of our LD-pruned estimator in Supplemental figures S3, S4, and S5. For the results from the LD-pruned estimator, we vary the number of SNPs within a window between 500 SNPs ($\approx$ 100KB), 5,000 SNPs ($\approx$ 1MB), and 50,000 SNPs ($\approx$ 10MB). We also ensure that our estimates are sensible by estimating the proportion of additive genetic variance from variants we infer to be shared. If the inference procedure works correctly, this number should be $1 - \hat{\rho}$. In Supplemental figure S7, we see that this is indeed the case.

We validated our result on BMI using an external cohort from the GIANT consortium [18]. This GWAS on BMI was performed on 718,734 individuals using an exome-targeted genotyping array. Using the LD-pruned estimator and a probability cutoff of 0.1, we found the proportion of SNP heritability from private variants is 0.522 (95% confidence interval: 0.390-0.653). This value is very close to the inferred value on the full data from the UK Biobank (0.492), giving us confidence that these results are robust across association studies.

Recent studies have highlighted the effects of stratification on polygenic risk scores [1, 16]. We wondered whether stratification could have an effect on our analyses. To test this, we repeated our analyses using only those SNPs showing stronger associations with the trait. Specifically, we employed $p$-value cutoffs, using only SNPs with a $p$-value lower than the cutoff (Figure S8). Broadly, we find two patterns: 1) for case-control association analyses, we see that decreasing the $p$-value results in estimates of the proportion of heritability from private variants approaching 1 and 2) for quantitative trait analyses, the proportion of the heritability attributable to private variants decreases.

There are two factors that contribute to these observations. First, for all traits analyzed, the power to detect an association will be lower for private variants than shared variants because private variants tend to have lower allele frequencies. Therefore, as the $p$-value cutoff decreases, we expect a lower proportion of heritability to come from private variants (as seen in the quantitative traits). Second, since the case-control associations are not balanced in sample size (i.e. $N_{cases} << N_{controls}$), the power to detect associations in this dataset will be low. The variants that are detected will tend to have large effects and will tend to be lower in frequency because these case-control associations are made up of cancer and disease phenotypes which are likely under negative selection. Thus, as the $p$-value cutoff becomes stricter, the remaining pool of associated variants tend to private.

[9] International HapMap Consortium, Kelly A. Frazer, Dennis G. Ballinger, David R. Cox, David A. Hinds, Laura L. Stuve, Richard A. Gibbs, John W. Belmont, Andrew Boudreau, Paul Hardenbol, Suzanne M. Leal, Shiran Pasternak, David A. Wheeler, Thomas D. Willis, Fuli Yu, Huanming Yang, Changqing Zeng, Yang Gao, Haoran Hu, Weitao Hu, Chaohua Li, Wei Lin, Siqi Liu, Hao Pan, Xiaoli Tang, Jian Wang, Wei Wang, Jun Yu, Bo Zhang, Qingrun Zhang, Hongbin Zhao, Hui Zhao, Jun Zhou, Stacey B. Gabriel, Rachel Barry, Brendan Blumenstiel, Amy Camargo, Matthew Defelice, Maura Faggart, Mary Goyette, Supriya Gupta, Jamie Moore, Huy Nguyen, Robert C. Onofrio, Melissa Parkin, Jessica Roy, Erich Stahl, Ellen Winchester, Liuda Ziaugra, David Altshuler, Yan Shen, Zhijian Yao, Wei Huang, Xun Chu, Yungang He, Li Jin, Yangfan Liu, Yayun Shen, Weiwei Sun, Haifeng Wang, Yi Wang, Ying Wang, Xiaoyan Xiong, Liang Xu, Mary M. Y. Waye, Stephen K. W. Tsui, Hong Xue, J. Tze-Fei Wong, Luana M. Galver, Jian-Bing Fan, Kevin Gunderson, Sarah S. Murray, Arnold R. Oliphant, Mark S. Chee, Alexandre Montpetit, Fanny Chagnon, Vincent Ferretti, Martin Leboeuf, Jean-François Olivier, Michael S. Phillips, Stéphanie Roumy, Clémentine Sallée, Andrei Verner, Thomas J. Hudson, Pui-Yan Kwok, Dongmei Cai, Daniel C. Koboldt, Raymond D. Miller, Ludmila Pawlikowska, Patricia Taillon-Miller, Ming Xiao, Lap-Chee Tsui, William Mak, You Qiang Song, Paul K. H. Tam, Yusuke Nakamura, Takahisa Kawaguchi, Takuya Kitamoto, Takashi Morizono, Atsushi Nagashima, Yozo Ohnishi, Akihiro Sekine, Toshihiro Tanaka, Tatsuhiko Tsunoda, Panos Deloukas, Christine P. Bird, Marcos Delgado, Emmanouil T. Dermitzakis, Rhian Gwilliam, Sarah Hunt, Jonathan Morrison, Don Powell, Barbara E. Stranger, Pamela Whittaker, David R. Bentley, Mark J. Daly, Paul I. W. de Bakker, Jeff Barrett, Yves R. Chretien, Julian Maller, Steve McCarroll, Nick Patterson, Itsik Pe'er, Alkes Price, Shaun Purcell, Daniel J. Richter, Pardis Sabeti, Richa Saxena, Stephen F. Schaffner, Pak C. Sham, Patrick Varilly, David Altshuler, Lincoln D. Stein, Lalitha Krishnan, Albert Vernon Smith, Marcela K. Tello-Ruiz, Gudmundur A. Thorisson, Aravinda Chakravarti, Peter E. Chen, David J. Cutler, Carl S. Kashuk, Shin Lin, Gonçalo R. Abecasis, Weihua Guan, Yun Li, Heather M. Munro, Zhaohui Steve Qin, Daryl J. Thomas, Gilean McVean, Adam Auton, Leonardo Bottolo, Niall Cardin, Susana Eyheramendy, Colin Freeman, Jonathan Marchini, Simon Myers, Chris Spencer, Matthew Stephens, Peter Donnelly, Lon R. Cardon, Geraldine Clarke, David M. Evans,

Andrew P. Morris, Bruce S. Weir, Tatsuhiko Tsunoda, James C. Mullikin, Stephen T. Sherry, Michael Feolo, Andrew Skol, Houcan Zhang, Changqing Zeng, Hui Zhao, Ichiro Matsuda, Yoshimitsu Fukushima, Darryl R. Macer, Eiko Suda, Charles N. Rotimi, Clement A. Adebamowo, Ike Ajayi, Toyin Aniagwu, Patricia A. Marshall, Chibuzor Nkwodimmah, Charmaine D. M. Royal, Mark F. Leppert, Missy Dixon, Andy Peiffer, Renzong Qiu, Alastair Kent, Kazuto Kato, Norio Niikawa, Isaac F. Adewole, Bartha M. Knoppers, Morris W. Foster, Ellen Wright Clayton, Jessica Watkin, Richard A. Gibbs, John W. Belmont, Donna Muzny, Lynne Nazareth, Erica Sodergren, George M. Weinstock, David A. Wheeler, Imtaz Yakub, Stacey B. Gabriel, Robert C. Onofrio, Daniel J. Richter, Liuda Ziaugra, Bruce W. Birren, Mark J. Daly, David Altshuler, Richard K. Wilson, Lucinda L. Fulton, Jane Rogers, John Burton, Nigel P. Carter, Christopher M. Clee, Mark Griffiths, Matthew C. Jones, Kirsten McLay, Robert W. Plumb, Mark T. Ross, Sarah K. Sims, David L. Willey, Zhu Chen, Hua Han, Le Kang, Martin Godbout, John C. Wallenburg, Paul L'Archevêque, Guy Bellemare, Koji Saeki, Hongguang Wang, Daochang An, Hongbo Fu, Qing Li, Zhen Wang, Renwu Wang, Arthur L. Holden, Lisa D. Brooks, Jean E. McEwen, Mark S. Guyer, Vivian Ota Wang, Jane L. Peterson, Michael Shi, Jack Spiegel, Lawrence M. Sung, Lynn F. Zacharia, Francis S. Collins, Karen Kennedy, Ruth Jamieson, and John Stewart. A second generation human haplotype map of over 3.1 million SNPs. *Nature*, 449(7164):851–861, October 2007.

[10] Jerome Kelleher, Alison M. Etheridge, and Gilean McVean. Efficient Coalescent Simulation and Genealogical Analysis for Large Sample Sizes. *PLOS Computational Biology*, 12(5):e1004842, May 2016.

[11] Bernard Y. Kim, Christian D. Huber, and Kirk E. Lohmueller. Inference of the Distribution of Selection Coefficients for New Nonsynonymous Mutations Using Large Samples. *Genetics*, page genetics.116.197145, January 2017.

[12] Hans R. Kunsch. The Jackknife and the Bootstrap for General Stationary Observations. *The Annals of Statistics*, 17(3):1217–1241, September 1989.

[13] Monkol Lek, Konrad J. Karczewski, Eric V. Minikel, Kaitlin E. Samocha, Eric Banks, Timothy Fennell, Anne H. O'Donnell-Luria, James S. Ware, Andrew J. Hill, Beryl B. Cummings, Taru Tukiainen, Daniel P. Birnbaum, Jack A. Kosmicki, Laramie E. Duncan, Karol Estrada, Fengmei Zhao, James Zou, Emma Pierce-Hoffman, Joanne Berghout, David N. Cooper, Nicole Deflaux, Mark DePristo, Ron Do, Jason Flannick, Menachem Fromer, Laura Gauthier, Jackie Goldstein, Namrata Gupta, Daniel Howrigan, Adam Kiezun, Mitja I. Kurki, Ami Levy Moonshine, Pradeep Natarajan, Lorena Orozco, Gina M. Peloso, Ryan Poplin, Manuel A. Rivas, Valentin Ruano-Rubio, Samuel A. Rose, Douglas M. Ruderfer, Khalid Shakir, Peter D. Stenson, Christine Stevens, Brett P. Thomas, Grace Tiao, Maria T. Tusie-Luna, Ben Weisburd, Hong-Hee Won, Dongmei Yu, David M. Altshuler, Diego Ardissino, Michael Boehnke, John Danesh, Stacey Donnelly, Roberto Elosua, Jose C. Florez, Stacey B. Gabriel, Gad Getz, Stephen J. Glatt, Christina M. Hultman, Sekar Kathiresan, Markku Laakso, Steven McCarroll, Mark I. McCarthy, Dermot McGovern, Ruth McPherson, Benjamin M. Neale, Aarno Palotie, Shaun M. Purcell, Danish Saleheen, Jeremiah M. Scharf, Pamela Sklar, Patrick F. Sullivan, Jaakko Tuomilehto, Ming T. Tsuang, Hugh C. Watkins, James G. Wilson, Mark J. Daly, Daniel G. MacArthur, and Exome Aggregation Consortium. Analysis of protein-coding genetic variation in 60,706 humans. *Nature*, 536(7616):285–291, August 2016.

[14] Huwenbo Shi, Gleb Kichaev, and Bogdan Pasaniuc. Contrasting the Genetic Architecture of 30 Complex Traits from Summary Association Data. *The American Journal of Human Genetics*, 99(1):139–153, July 2016.

[15] Montgomery Slatkin. Linkage disequilibrium–understanding the evolutionary past and mapping the medical future. *Nature Reviews. Genetics*, 9(6):477–485, June 2008.

[16] Mashaal Sohail, Robert M Maier, Andrea Ganna, Alex Bloemendal, Alicia R Martin, Michael C Turchin, Charleston WK Chiang, Joel Hirschhorn, Mark J Daly, Nick Patterson, Benjamin Neale, Iain Mathieson, David Reich, and Shamil R Sunyaev. Polygenic adaptation on height is overestimated due to uncorrected stratification in genome-wide association studies. *eLife*, 8:e39702, March 2019.

[17] Doug Speed and David J. Balding. SumHer better estimates the SNP heritability of complex traits from summary statistics. *Nature Genetics*, 51(2):277, February 2019.

[18] Valérie Turcot, Yingchang Lu, Heather M. Highland, Claudia Schurmann, Anne E. Justice, Rebecca S. Fine, Jonathan P. Bradfield, Tõnu Esko, Ayush Giri, Mariaelisa Graff, Xiuqing Guo, Audrey E. Hendricks, Tugce Karaderi, Adelheid Lempradl, Adam E. Locke, Anubha Mahajan, Eirini Marouli, Suthesh Sivapalaratnam, Kristin L. Young, Tamuno Alfred, Mary F. Feitosa, Nicholas G. D. Masca, Alisa K. Manning, Carolina Medina-Gomez, Poorva Mudgal, Maggie C. Y. Ng, Alex P. Reiner, Sailaja Vedantam, Sara M. Willems, Thomas W. Winkler, Gonçalo Abecasis, Katja K. Aben, Dewan S. Alam, Sameer E. Alharthi, Matthew Allison, Philippe Amouyel, Folkert W. Asselbergs, Paul L. Auer, Beverley Balkau, Lia E. Bang, Inês Barroso, Lisa Bastarache, Marianne Benn, Sven Bergmann, Lawrence F. Bielak, Matthias Blüher, Michael Boehnke, Heiner Boeing, Eric Boerwinkle, Carsten A. Böger, Jette Bork-Jensen, Michiel L. Bots, Erwin P. Bottinger, Donald W. Bowden, Ivan Brandslund, Gerome Breen, Murray H. Brilliant, Linda Broer, Marco Brumat, Amber A. Burt, Adam S. Butterworth, Peter T. Campbell, Stefania Cappellani, David J. Carey, Eulalia Catamo, Mark J. Caulfield, John C. Chambers, Daniel I. Chasman, Yii-Der I. Chen, Rajiv Chowdhury, Cramer Christensen, Audrey Y. Chu, Massimiliano Cocca, Francis S. Collins, James P. Cook, Janie Corley, Jordi Corominas Galbany, Amanda J. Cox, David S. Crosslin, Gabriel Cuellar-Partida, Angela D'Eustacchio, John Danesh, Gail Davies, Paul I. W. Bakker, Mark C. H. Groot, Renée Mutsert, Ian J. Deary, George Dedoussis, Ellen W. Demerath, Martin Heijer, Anneke I. Hollander, Hester M. Ruijter, Joe G. Dennis, Josh C. Denny, Emanuele Di Angelantonio, Fotios Drenos, Mengmeng Du, Marie-Pierre Dubé, Alison M. Dunning, Douglas F. Easton, Todd L. Edwards, David Ellinghaus, Patrick T. Ellinor, Paul Elliott, Evangelos Evangelou, Aliki-Eleni Farmaki, I. Sadaf Farooqi, Jessica D. Faul, Sascha Fauser, Shuang Feng, Ele Ferrannini, Jean Ferrieres, Jose C. Florez, Ian Ford, Myriam Fornage, Oscar H. Franco, Andre Franke, Paul W. Franks, Nele Friedrich, Ruth Frikke-Schmidt, Tessel E. Galesloot, Wei Gan, Ilaria Gandin, Paolo Gasparini, Jane Gibson, Vilmantas Giedraitis, Anette P. Gjesing, Penny Gordon-Larsen, Mathias Gorski, Hans-Jörgen Grabe, Struan F. A. Grant, Niels Grarup, Helen L. Griffiths, Megan L. Grove, Vilmundur Gudnason, Stefan Gustafsson, Jeff Haessler, Hakon Hakonarson, Anke R. Hammerschlag, Torben Hansen, Kathleen Mullan Harris, Tamara B. Harris, Andrew T. Hattersley, Christian T. Have, Caroline Hayward, Liang He, Nancy L. Heard-Costa, Andrew C. Heath, Iris M. Heid, Øyvind Helgeland, Jussi Hernesniemi, Alex W. Hewitt, Oddgeir L. Holmen, G. Kees Hovingh, Joanna M. M. Howson, Yao Hu, Paul L. Huang, Jennifer E. Huffman, M. Arfan Ikram,

Erik Ingelsson, Anne U. Jackson, Jan-Håkan Jansson, Gail P. Jarvik, Gorm B. Jensen, Yucheng Jia, Stefan Johansson, Marit E. Jørgensen, Torben Jørgensen, J. Wouter Jukema, Bratati Kahali, René S. Kahn, Mika Kähönen, Pia R. Kamstrup, Stavroula Kanoni, Jaakko Kaprio, Maria Karaleftheri, Sharon L. R. Kardia, Fredrik Karpe, Sekar Kathiresan, Frank Kee, Lambertus A. Kiemeney, Eric Kim, Hidetoshi Kitajima, Pirjo Komulainen, Jaspal S. Kooner, Charles Kooperberg, Tellervo Korhonen, Peter Kovacs, Helena Kuivaniemi, Zoltán Kutalik, Kari Kuulasmaa, Johanna Kuusisto, Markku Laakso, Timo A. Lakka, David Lamparter, Ethan M. Lange, Leslie A. Lange, Claudia Langenberg, Eric B. Larson, Nanette R. Lee, Terho Lehtimäki, Cora E. Lewis, Huaixing Li, Jin Li, Ruifang Li-Gao, Honghuang Lin, Keng-Hung Lin, Li-An Lin, Xu Lin, Lars Lind, Jaana Lindström, Allan Linneberg, Ching-Ti Liu, Dajiang J. Liu, Yongmei Liu, Ken S. Lo, Artitaya Lophatananon, Andrew J. Lotery, Anu Loukola, Jian'an Luan, Steven A. Lubitz, Leo-Pekka Lyytikäinen, Satu Männistö, Gaëlle Marenne, Angela L. Mazul, Mark I. McCarthy, Roberta McKean-Cowdin, Sarah E. Medland, Karina Meidtner, Lili Milani, Vanisha Mistry, Paul Mitchell, Karen L. Mohlke, Leena Moilanen, Marie Moitry, Grant W. Montgomery, Dennis O. Mook-Kanamori, Carmel Moore, Trevor A. Mori, Andrew D. Morris, Andrew P. Morris, Martina Müller-Nurasyid, Patricia B. Munroe, Mike A. Nalls, Narisu Narisu, Christopher P. Nelson, Matt Neville, Sune F. Nielsen, Kjell Nikus, Pål R. Njølstad, Børge G. Nordestgaard, Dale R. Nyholt, Jeffrey R. O'Connel, Michelle L. O'Donoghue, Loes M. Olde Loohuis, Roel A. Ophoff, Katharine R. Owen, Chris J. Packard, Sandosh Padmanabhan, Colin N. A. Palmer, Nicholette D. Palmer, Gerard Pasterkamp, Aniruddh P. Patel, Alison Pattie, Oluf Pedersen, Peggy L. Peissig, Gina M. Peloso, Craig E. Pennell, Markus Perola, James A. Perry, John R. B. Perry, Tune H. Pers, Thomas N. Person, Annette Peters, Eva R. B. Petersen, Patricia A. Peyser, Ailith Pirie, Ozren Polasek, Tinca J. Polderman, Hannu Puolijoki, Olli T. Raitakari, Asif Rasheed, Rainer Rauramaa, Dermot F. Reilly, Frida Renström, Myriam Rheinberger, Paul M. Ridker, John D. Rioux, Manuel A. Rivas, David J. Roberts, Neil R. Robertson, Antonietta Robino, Olov Rolandsson, Igor Rudan, Katherine S. Ruth, Danish Saleheen, Veikko Salomaa, Nilesh J. Samani, Yadav Sapkota, Naveed Sattar, Robert E. Schoen, Pamela J. Schreiner, Matthias B. Schulze, Robert A. Scott, Marcelo P. Segura-Lepe, Svati H. Shah, Wayne H.-H. Sheu, Xueling Sim, Andrew J. Slater, Kerrin S. Small, Albert V. Smith, Lorraine Southam, Timothy D. Spector, Elizabeth K. Speliotes, John M. Starr, Kari Stefansson, Valgerdur Steinthorsdottir, Kathleen E. Stirrups, Konstantin Strauch, Heather M. Stringham, Michael Stumvoll, Liang Sun, Praveen Surendran, Amy J. Swift, Hayato Tada, Katherine E. Tansey, Jean-Claude Tardif, Kent D. Taylor, Alexander Teumer, Deborah J. Thompson, Gudmar Thorleifsson, Unnur Thorsteinsdottir, Betina H. Thuesen, Anke Tönjes, Gerard Tromp, Stella Trompet, Emmanouil Tsafantakis, Jaakko Tuomilehto, Anne Tybjaerg-Hansen, Jonathan P. Tyrer, Rudolf Uher, André G. Uitterlinden, Matti Uusitupa, Sander W. Laan, Cornelia M. Duijn, Nienke Leeuwen, Jessica van Setten, Mauno Vanhala, Anette Varbo, Tibor V. Varga, Rohit Varma, Digna R. Velez Edwards, Sita H. Vermeulen, Giovanni Veronesi, Henrik Vestergaard, Veronique Vitart, Thomas F. Vogt, Uwe Völker, Dragana Vuckovic, Lynne E. Wagenknecht, Mark Walker, Lars Wallentin, Feijie Wang, Carol A. Wang, Shuai Wang, Yiqin Wang, Erin B. Ware, Nicholas J. Wareham, Helen R. Warren, Dawn M. Waterworth, Jennifer Wessel, Harvey D. White, Cristen J. Willer, James G. Wilson, Daniel R. Witte, Andrew R. Wood, Ying Wu, Hanieh Yaghootkar, Jie Yao, Pang Yao, Laura M. Yerges-Armstrong, Robin Young, Eleftheria Zeggini, Xiaowei Zhan, Weihua Zhang, Jing Hua Zhao, Wei Zhao, Wei

Zhao, Wei Zhou, Krina T. Zondervan, Jerome I. Rotter, John A. Pospisilik, Fernando Rivadeneira, Ingrid B. Borecki, Panos Deloukas, Timothy M. Frayling, Guillaume Lettre, Kari E. North, Cecilia M. Lindgren, Joel N. Hirschhorn, Ruth J. F. Loos, CHD Exome+ Consortium, EPIC-CVD Consortium, ExomeBP Consortium, Global Lipids Genetic Consortium, GoT2D Genes Consortium, EPIC InterAct Consortium, INTERVAL Study, ReproGen Consortium, T2D-Genes Consortium, MAGIC Investigators, and Understanding Society Scientific Group. Protein-altering variants associated with body mass index implicate pathways that control energy intake and expenditure in obesity. *Nature Genetics*, 50(1):26–41, 2018.

[19] Jenna M. VanLiere and Noah A. Rosenberg. Mathematical properties of the r2 measure of linkage disequilibrium. *Theoretical Population Biology*, 74(1):130–137, August 2008.

[20] J. Wakeley and J. Hey. Estimating ancestral population parameters. *Genetics*, 145(3):847–855, March 1997.

| AFR frequency cutoff | 0 | 0.001 | 0.005 | 0.01 | 0.05 |
|---|---|---|---|---|---|
| Predicted private, not observed in AFR | 0.83 | 0.93 | 0.95 | 0.96 | 0.97 |
| Predicted private, observed in AFR | 0.15 | 0.06 | 0.04 | 0.03 | 0.01 |
| Not predicted private, observed in AFR | 0 | 0 | 0 | 0 | 0 |
| Not predicted private, not observed in AFR | 0.015 | 0.015 | 0.015 | 0.015 | 0.015 |

Table S1: Empirical validation of private allele model with allele frequency filters. We tested the ability of our model (See Section S3) to correctly call alleles private in a sample of 66,740 chromosomes from Europe and 10,406 chromosomes from a mixture of African and African-American samples from the Exome Aggregation Consotrium (ExAC) with a posterior probability cutoff of 0.23. Because private European alleles can occur at a low frequency in the African-American samples due to admixture, we use a frequency filter in the AFR sample to remove these false shared variants.

| Threshold | Sample set | FDR | FNR | FPR | TPR |
|---|---|---|---|---|---|
| 0.1 | 10K Simulated | 0.062 | 0.005 | 0.483 | 0.995 |
| 0.1 | 50K Simulated | 0.044 | 0.001 | 0.692 | 0.999 |
| 0.1 | ExAC NFE | 0.144 | 0.001 | 0.908 | 0.999 |
| 0.23 | 10K Simulated | 0.555 | 0.017 | 0.422 | 0.983 |
| 0.23 | 50K Simulated | 0.04 | 0.004 | 0.636 | 0.996 |
| 0.23 | ExAC NFE | 0.141 | 0.003 | 0.884 | 0.997 |
| 0.4 | 10K Simulated | 0.049 | 0.047 | 0.363 | 0.952 |
| 0.4 | 50K Simulated | 0.039 | 0.011 | 0.608 | 0.989 |
| 0.4 | ExAC NFE | 0.138 | 0.005 | 0.865 | 0.995 |

Table S2: Simulated and empirical validation of the private allele model across different posterior probability thresholds (first column). For the results presented in the main text, we use a posterior probability cutoff of 0.23, which results in a simulated FDR of $\approx 5\%$ and an empirical FDR of $\approx 14\%$. We use the 0.1 threshold in Figures S3, S4, S5, S6, S7, S8. "ExAC NFE" refers to an empirical test from the ExAC data set where we use the observed frequency of the allele in the African American sample to estimate the allele count in a sample of unadmixed African genomes (see Section S3).

| $\tau$ | Shared (Europe) | Private (Europe) | Shared (Africa) | Private (Africa) |
|---|---|---|---|---|
| 0 | 0.91 | 0.38 | 0.96 | 0.34 |
| 0.25 | 0.76 | 0.62 | 0.83 | 0.57 |
| 0.5 | 0.56 | 0.83 | 0.58 | 0.79 |

Table S3: Correlation coefficient (Pearson's $r^2$) between the true genetic risk and the PRS using shared or private variation in European and African individuals across values of $\tau$ ($p < 2 \times 10^{-16}$ for all correlations). As negative selection becomes stronger, the correlation between the true genetic risk and shared variants decreases, while the correlation between true genetic risk and private variation increases.

| $\tau$ | $C$ | $h^2$ |
|---|---|---|
| 0 | 0.1 | 0.38 |
| 0.25 | 0.6 | 0.41 |
| 0.5 | 1.8 | 0.39 |

Table S4: Values of $C$ used in simulations for varying values of $\tau$ and the average resulting heritability. Values of $C$ were chosen to obtain $h^2 \approx 0.4$.

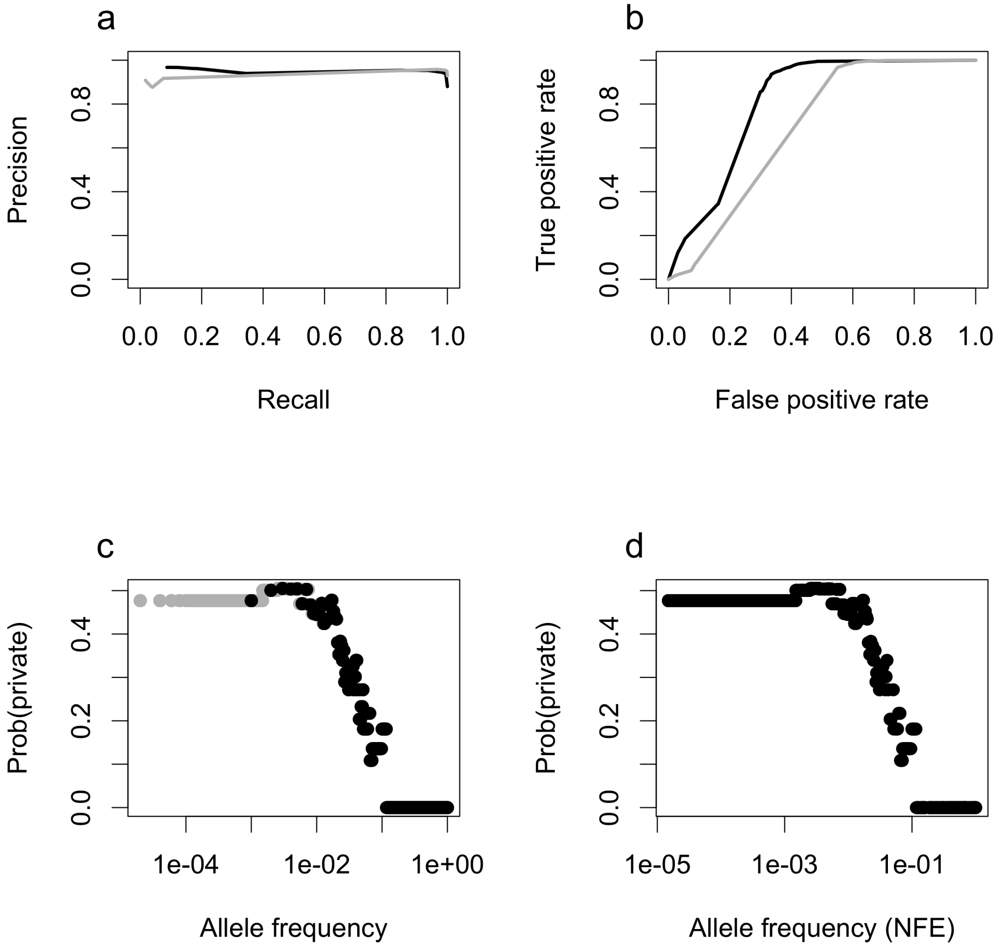

Figure S1: Performance and external validation of our model for inferring whether a variant is private or shared. a) precision-recall and b) receiver operating characteristic curve. c) The probability that a variant is private versus the allele frequency (log scale) in simulated test data. We see that for variants above 10%, the probability they are private is very low. In panels a,b, and c, the grey lines and points indicate results from a simulation with 50,000 haplotypes. d) The probability that a variant is private versus the allele frequency (log scale) for 10,000 randomly sampled variants from ExAC. Variants with a frequency below $1 \times 10^{-3}$ automatically set to have a probability equal to a variant at frequency 0.001 since these variants are outside the frequency range of our simulations.

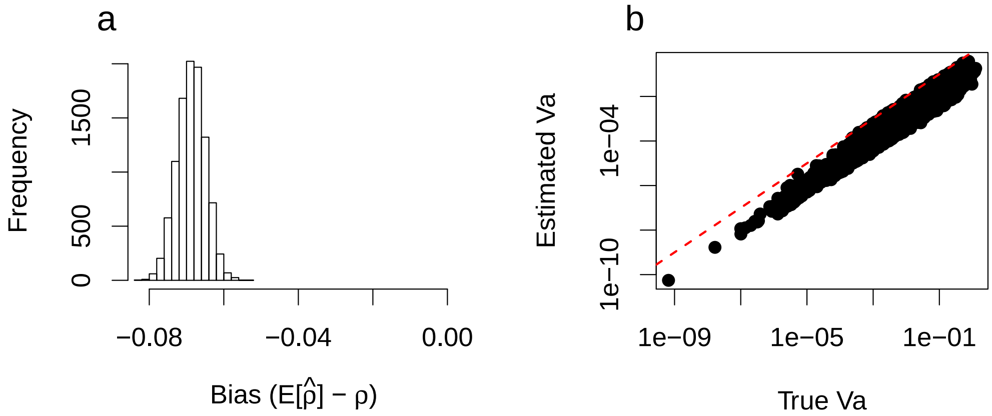

Figure S2: Bias of the LD-pruned estimator based on coalescent simulations a) We plot the expected value of $\hat{\rho}$ versus the true value of $\rho$. Using 10,000 simulations, we compute $E[\hat{\rho}]$ as well as $\rho$ and plot the difference. The distribution represents bootstrap re-sampled values across simulations. We see that the estimate of the proportion of $V_a$ from private variants is underestimated by $6-8\%$. b) Estimated $V_a$ versus the true $V_a$ from simulations for private variants. The red dashed line represents the 1:1 line. All points fall below this line, again suggesting this inference is downwardly biased.

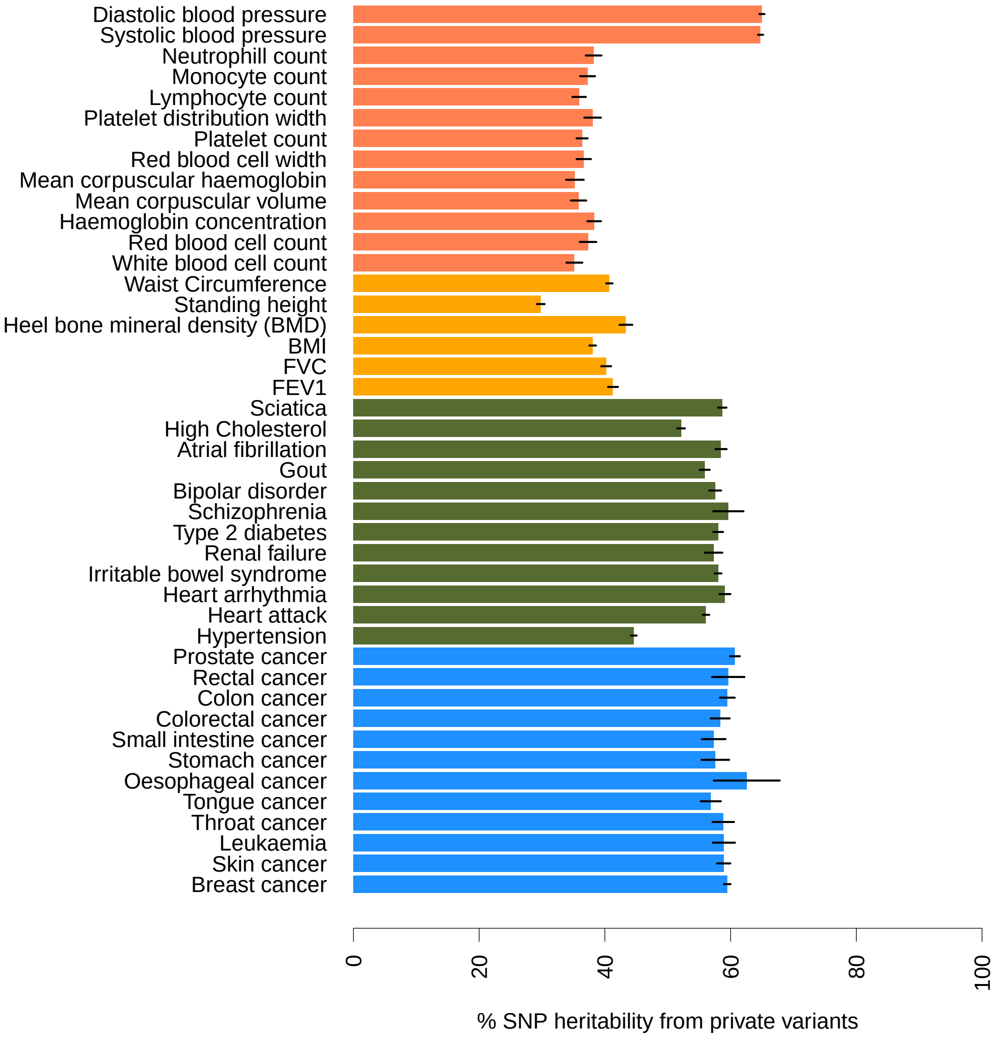

Figure S3: LD pruned estimator for the percentage of heritability from private variants. We use 500 SNPs per window. Lines indicate 95% confidence intervals obtained via a jackknife over the windows.

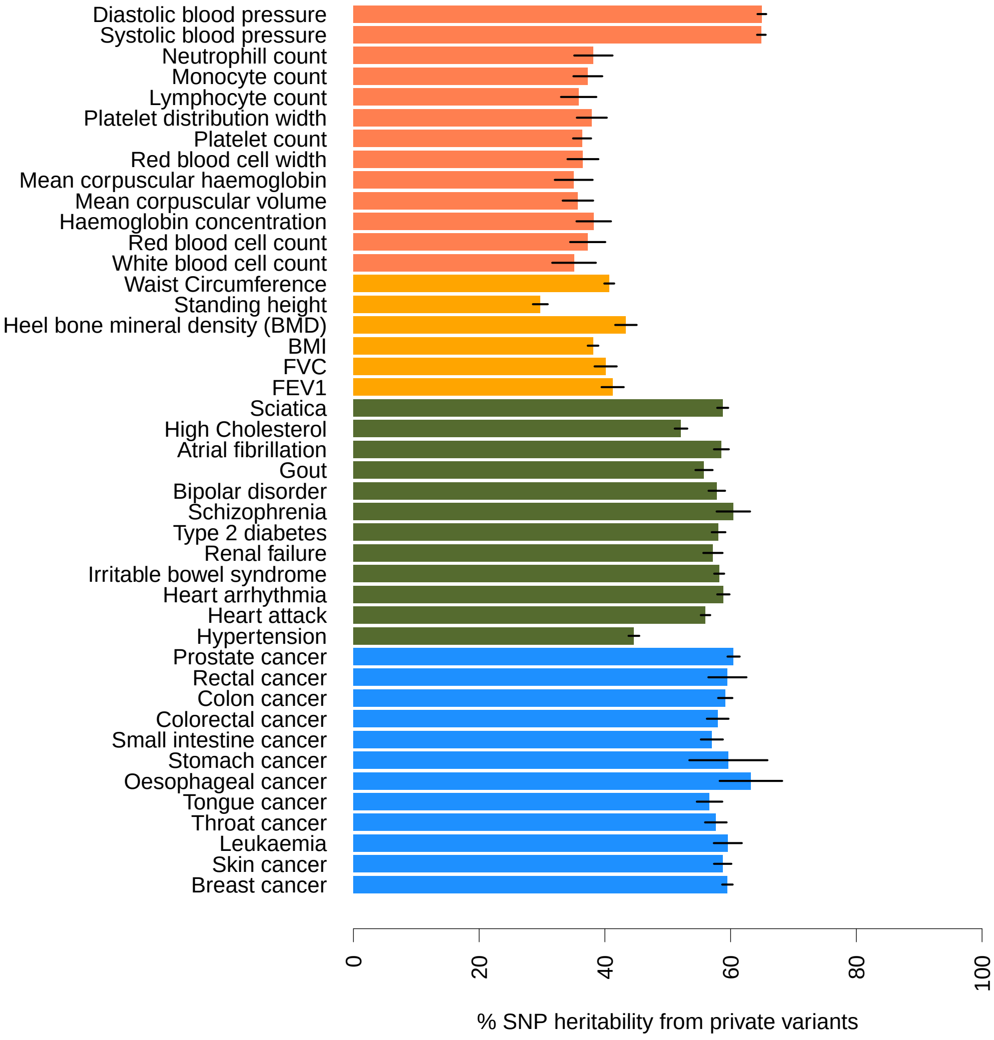

Figure S4: LD pruned estimator for the percentage of heritability from private variants. We use 5,000 SNPs per window. Lines indicate 95% confidence intervals obtained via a jackknife over the windows.

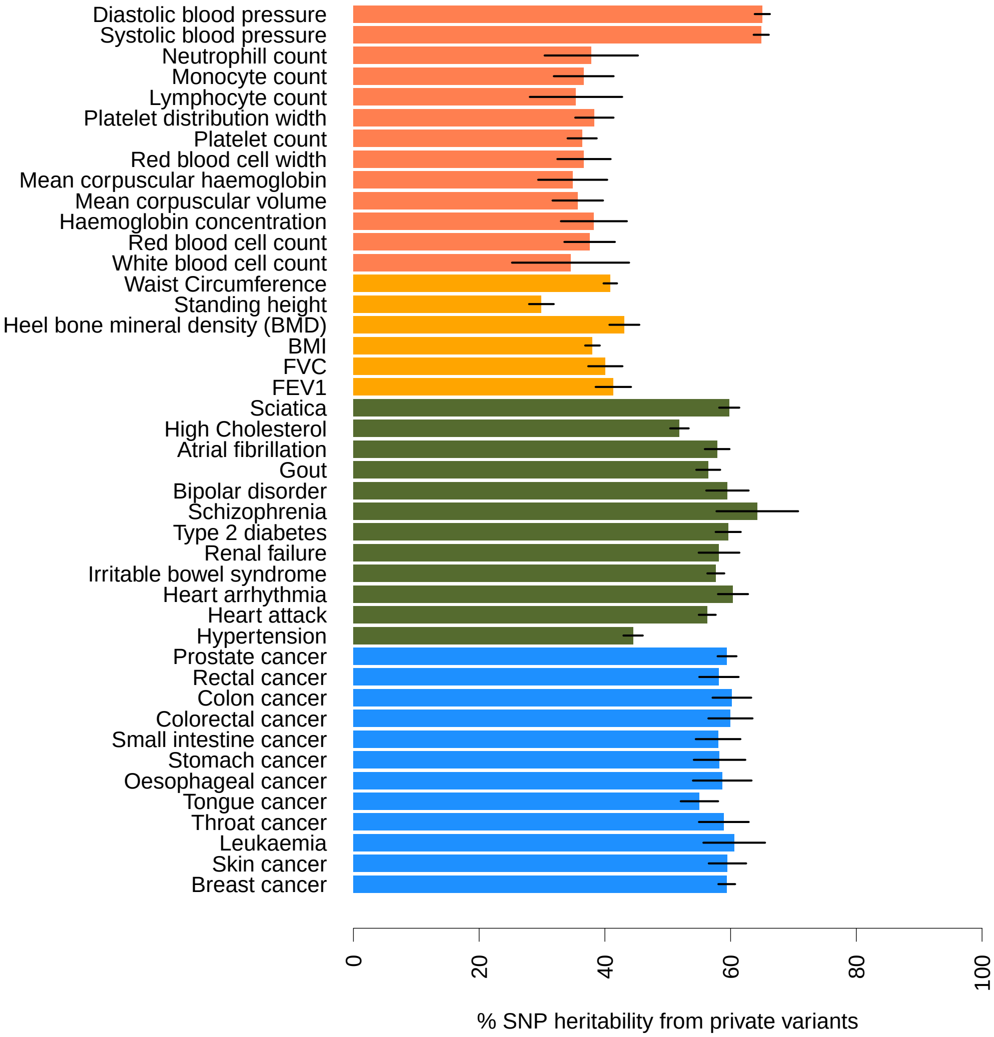

Figure S5: LD pruned estimator for the percentage of heritability from private variants. We use 50,000 SNPs per window. Lines indicate 95% confidence intervals obtained via a jackknife over the windows.

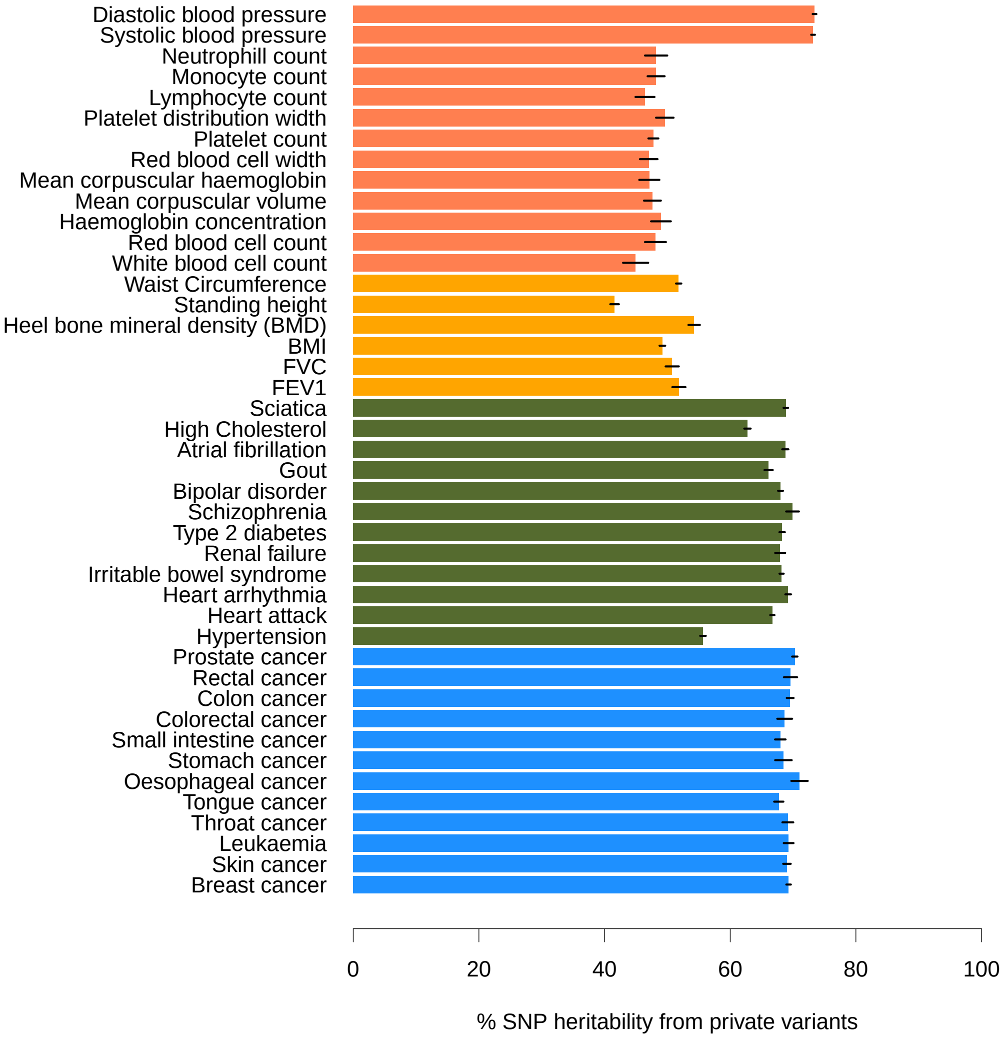

Figure S6: Fraction of heritability explained by private variants across 43 traits and diseases in the UK Biobank using a posterior probability cutoff of 0.1 for variants being called private. Lines indicate 95% confidence intervals obtained via a jackknife over the windows.

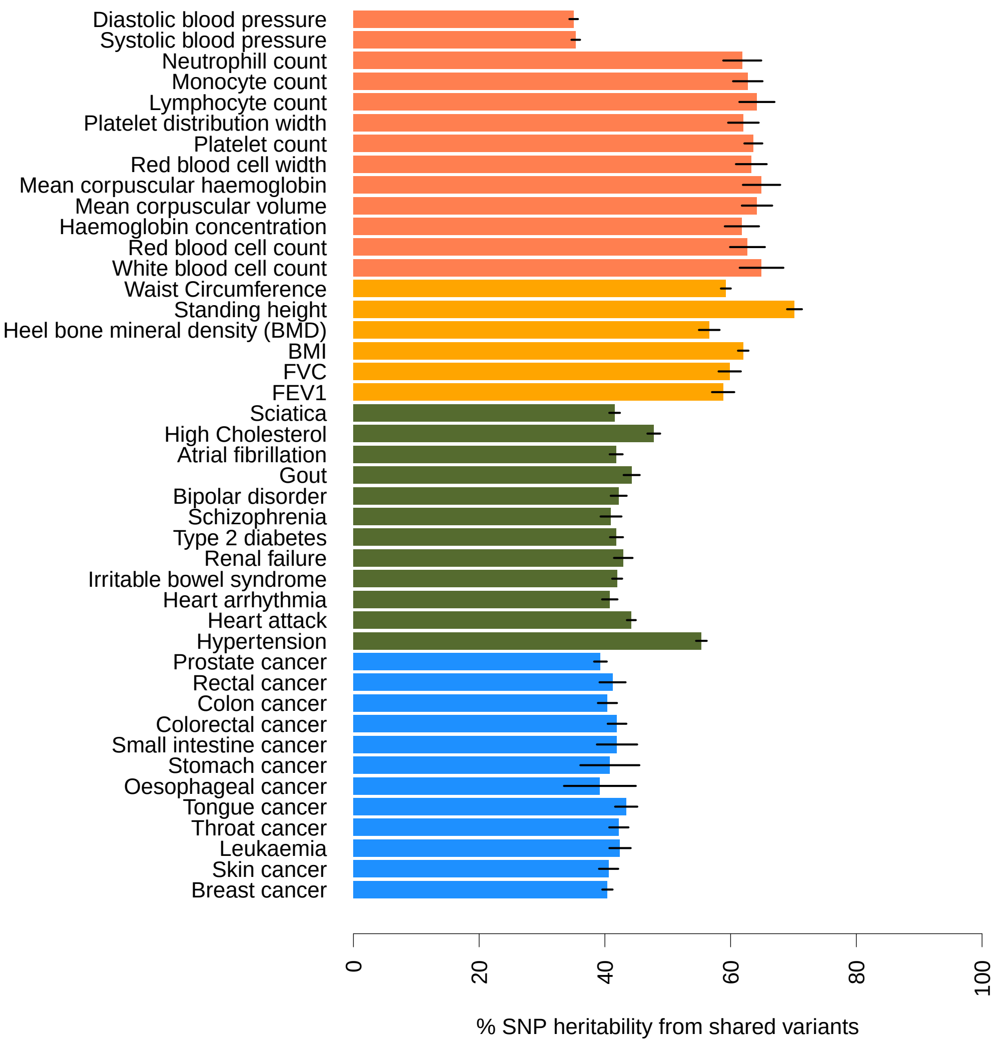

Figure S7: LD pruned estimator for the percentage of heritability from shared variants. We use 5,000 SNPs per window. Lines indicate 95% confidence intervals obtained via a jackknife over the windows.

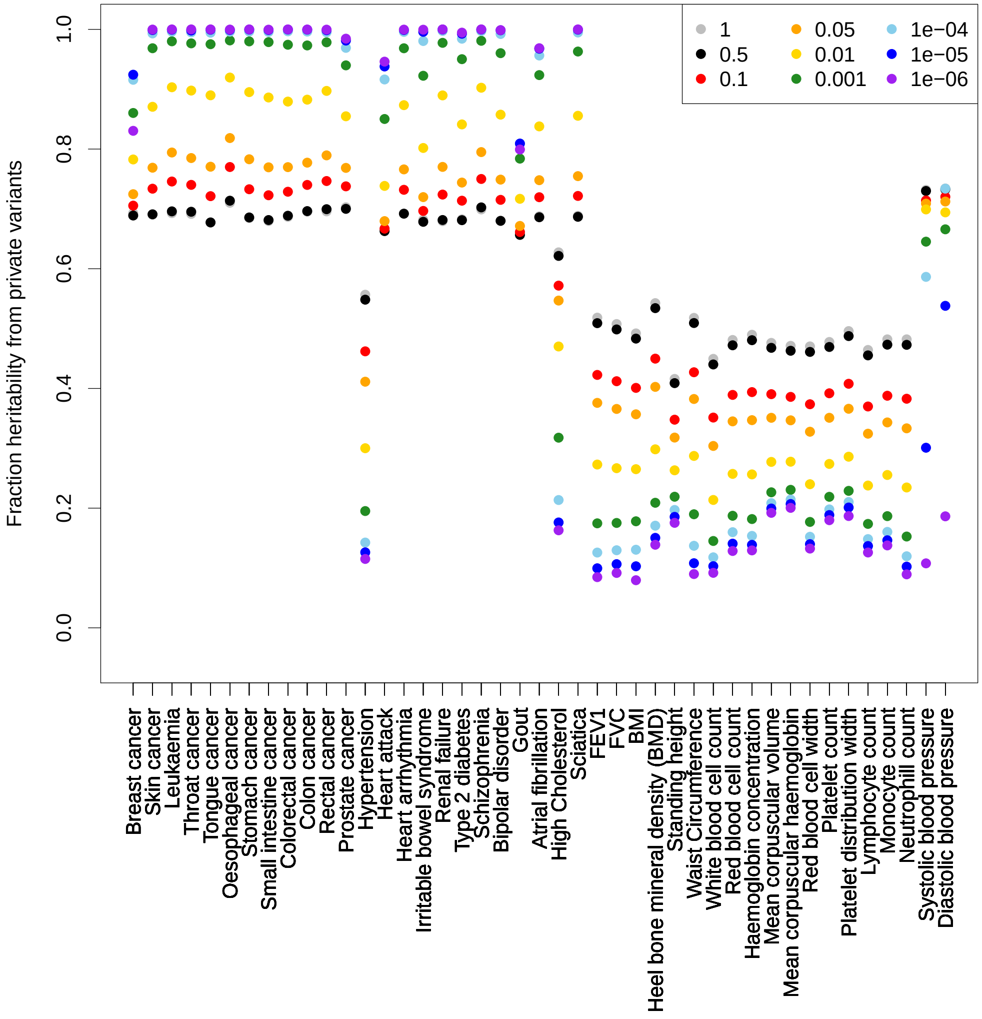

Figure S8: The fraction of heritability explained by private variants. For each trait, we calculated this fraction using a $p$-value cutoff (indicated by color). They grey dots represent no filter.
